# A Multitrait Genome-Wide Association Study Reveals a Requirement for the Strigolactone Receptor MtDWARF14 in Optimal GOLVEN Signaling

**DOI:** 10.1101/2024.06.24.599968

**Authors:** Sonali Roy, Yun Kang, Shulan Zhang, Ivone Torres-Jerez, Divya Jain, Bailey Sanchez, Liana Burghardt, Xiaofei Cheng, Jiangqi Wen, Jeremy D. Murray, Wolf-Rüdiger Scheible, Michael Udvardi

## Abstract

GOLVEN/ROOT MERISTEM GROWTH FACTOR family of signaling peptides have been shown to control root lateral organ number, density and positioning in plants, although the signaling pathways involved remain obscure. A diverse set of 171 *Medicago truncatula* HapMap accessions with variation in responses to the GOLVEN 10 peptide, GLV10, were used to identify 74 significant loci controlling seven traits related to nodule formation and root architecture. Importantly, a single nucleotide polymorphism (SNP) in the upstream region of the MtGLV10 peptide-inducible strigolactone receptor gene, *MtDWARF14* was significantly associated with insensitivity of nodule density to GLV10, suggesting a link between strigolactone signaling and GLV10 responsiveness. Three independent *d14* mutants of the D*WARF14* gene were found to hypernodulate, while overexpression of the gene led to reduction in nodule number, phenocopying GLV10. A null mutant, *mtd14-1*, remained sensitive to GLV10’s effect on nodule density. However, at the transcriptional level, the mutant failed to effectively induce the expression of the GOLVEN marker genes, *MtPLETHORA3* and *MtPINLIKES2*. Our study uncovers a hitherto unknown link between the strigolactone and GLV peptide signaling pathways using genotype x environment analysis of Medicago HapMap lines and provides a putative molecular mechanism for recovery from frost damage to fine roots.

## INTRODUCTION

Poor nutrient availability, especially of macronutrients like Nitrogen (N) and Phosphorus (P), limit plant growth and consequently food production (Udvardi et al., 2021). Plants regulate root development and architecture to optimize acquisition of soil nutrients, which are heterogeneously distributed in soil and vary in concentration over time and space (Mohd-Radzman et al., 2013; Rellán-Álvarez et al., 2016). Nutrient acquisition involves regulation of primary and secondary (lateral) root formation and, in the case of nitrogen-fixing plants like legumes, initiation of specialized lateral root organs called nodules that accommodate nitrogen-fixing bacteria called rhizobia (Roy et al., 2020). This ability of legumes to fix atmospheric nitrogen in cooperation with soil bacteria helps reduce the need for synthetic fertilizers in agriculture. With increasing unpredictability in weather patterns that alter legume cultivation globally, optimizing root and nodule development to sustain or enhance crop yields remains a high agricultural priority (Udvardi et al., 2021; Yu et al., 2021). To date, studies on model legumes such as *Medicago truncatula* have helped identify genes that are required for nodule formation, many of which are shared with those required for lateral root formation (Hirsch et al., 1997; Roy et al., 2020). Indeed, nodules are considered modified lateral roots with a significant ontological and genetic overlap between the two developmental programs (Herrbach et al., 2014; Xiao et al., 2014; Schiessl et al., 2019).

Several external factors such as light, and internal signals such as hormones control root and nodule development in response to nutrient supply and plant demand (Roy et al., 2020; Ji et al., 2022; Lee et al., 2024). Of note are small signaling peptides or peptide hormones which are short chains of amino acids that are biologically active in modulating plant physiology because they are perceived by cell surface receptors that then trigger downstream signaling cascades (Roy and Muller, 2022). Peptide hormones can act locally or systemically to control lateral root and nodule initiation and development (Tabata et al., 2014). One such peptide family is the GOLVEN (GLV)/ROOT MERISTEM GROWTH FACTOR (RGF). The sulfated GLV/RGF family of peptides plays an important role in nodulation controlling nodule number, density, noduletaxis, nodule zone length and, several root architecture traits including primary root length, lateral root density (Fernandez et al., 2015; Fernandez et al., 2020; Li et al., 2020; Jourquin et al., 2022; Roy et al., 2024). Importantly, GOLVEN peptide-coding genes transcriptionally regulated by auxin, and in turn, modulate both nodulation and root architectural traits through interference with auxin accumulation and signaling pathways, pointing at the existence of a feedback loop (Whitford et al., 2012; Jourquin et al., 2023; Roy et al., 2024). Approximately 30% of genes controlled by GLV10 transcriptionally are also regulated by auxin, including auxin efflux transporters PINs and PIN-LIKEs (Roy et al., 2024). In Arabidopsis, the AtGLV6 peptide affects the turnover of PIN2 transporters in the plasma membrane of the root epidermis thereby dissipating auxin maxima required for subsequent lateral root formation (Whitford et al., 2012). Since legumes not only interact with beneficial rhizobia to acquire nitrogen but can also modulate their root architecture to absorb nitrogen from the soil, GLVs are critical for optimal nitrogen acquisition (Mohd-Radzman et al., 2013). Further research is required to uncover additional components of the signaling pathway(s) that mediate responses to GLV peptides, since knowledge of these may help to optimize root and nodule development for nutrient acquisition in plants.

Over the last two decades, artificially induced genetic variation created using insertional mutagens or interbreeding has developed populations that have revealed more than 200 genes essential for the legume-rhizobia symbiosis and other developmental traits (Thoquet et al., 2002; Tadege et al., 2008; Pislariu et al., 2012; Małolepszy et al., 2016; Roy et al., 2020). Genome Wide Association Studies (GWAS) is a newer experimental approach that uses natural genomic variations correlated with observed phenotypic variations in a population, using statistical methods to identify novel determinants underlying a trait (Uffelmann et al., 2021). Resources in *M. truncatula* that make it a model plant to study root nodule symbiosis include a robust sequenced population of Haplotype mapping or ‘HapMap’ lines with sufficient genomic sequence variation to conduct GWA studies (Nandety et al., 2023). GWAS using the Medicago HapMap lines have helped uncover genomic loci that influence the phenotype of interest such as climatic adaptation, salt stress, seed size and composition, disease resistance and, ability to form nodules (Rey et al., 2017; Guerrero et al., 2018; Kang et al., 2019; Bonhomme et al., 2021; Chen et al., 2021a; Chen et al., 2021b). However, phenotyping below-ground traits such as roots remains a significant technical challenge in plant-microbe interactions and therefore accounts for less than 15% of GWAS conducted worldwide (Demirjian et al., 2023). Additionally, prioritizing causal SNPs is an analytical problem but integrating GWAS with expression data has proven successful in narrowing down gene candidates (Schaefer et al., 2018). Despite these challenges, previous GWAS on nodulation have identified several known genes that were initially discovered through reverse genetics and other approaches, validating this method as a robust approach for identifying genomic loci associated with complex traits like root nodule symbiosis (Stanton-Geddes et al., 2013; Epstein et al., 2023). These differences which manifest as single nucleotide polymorphisms (SNPs), often a result of ecological and environmental pressures, offer insights into how plants adapt to diverse climatic and biogeographical conditions (Guerrero et al., 2018; Epstein et al., 2023).

We hypothesized that variation in response to GLV10 exists within the *M. truncatula* species and can be leveraged to identify genes involved in GLV10 signaling, via GWAS. Here, we verify this hypothesis and identify numerous genetic loci associated with variation in response to the peptide GLV10, including a gene involved in strigolactone perception, *MtDWARF14*. The results are important as they show a connection between GLV and strigolactone signaling, as well as offering additional leads to genes potentially involved in GLV control of root and nodule development.

## RESULTS

### Analysis of Natural Variation in GLV10 peptide Sensitivity Among *M. truncatula* Accessions

To understand how GLV peptides bring about their effects on root nodulation and architecture, we leveraged natural variation among *M. truncatula* HapMap accessions and conducted GWAS of GLV10 peptide (GLV10p) sensitivity. Among the fifteen GOLVEN peptides encoded by *M. truncatula*, GLV10 consistently exhibited the most pronounced and reliable effects on root growth and nodulation traits (Roy et al., 2024). Additionally, the visually evident agravitropic response served as a clear indicator of the treatment’s efficacy. Consequently, we selected this peptide for our GWAS study (**Figure 1A**).

**Figure 1.**
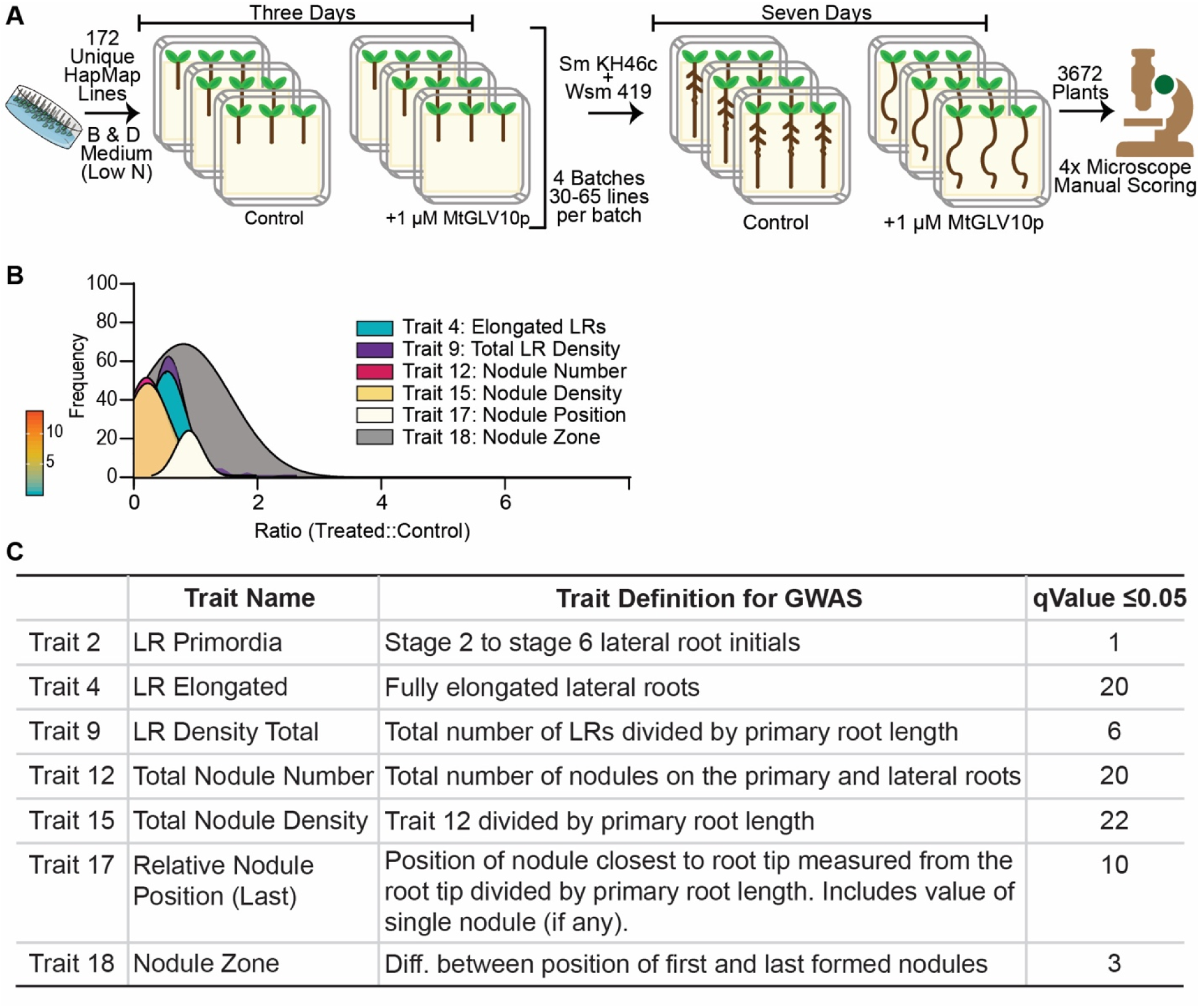
Overview of GWAS study and effect of GLV10 peptide treatment on traits assessed in this study. **A.** Diagrammatic overview of the GWAS setup using MtGLV10p **B.** Plot showing data distribution of root traits with significant SNPs using histogram curve. **C.** Table showing root traits with significant SNPs with a q value ≤0.05.

A population of 171 lines of an existing 227 Medicago population were selected after analyzing maximum diversity based on kinship and the most genetically diverse lines were prioritized using the R function “CoreSetter” with 8000 randomly selected SNPs **(Supplementary Table 1)**. These lines represent a collection of *M. truncatula* accessions from the Mediterranean region that also include a few lines from North America (Yoder et al., 2014). We screened these 171 lines under highly controlled, sterile conditions using the filter-paper sandwich system for Medicago growth (Stanton-Geddes et al., 2013; Boschiero et al., 2019). To overcome feasibility issues of handling many lines with sufficient replicates, we split the selected HapMap population into four batches with ∼30-60 lines per batch **(Supplementary Table 1)**. Each batch was infected with a mixture of two rhizobial strains, *Sinorhizobium meliloti* KH46c and *Sinorhizobium medicae* WSM419, to ameliorate any differences in strain compatibility with different plant genotypes (**Figure 1A**). To avoid block effects, nine seedlings were grown on three different agarose plates for both control and treated plants under controlled conditions of plant growth (See materials and methods for more details). All primary traits i.e those not dependent on another root growth parameter (e.g. - Root length, number of lateral organs, nodule position) were scored manually under the microscope and the derived traits (lateral root density, relative nodule position) were calculated relative to the root length or nod zone. To measure the effect of GLV10p, we normalized the treated values by dividing them by the untreated (control) baseline values. Since the peptide treatment led to an organ count of zero for several traits such as root nodules and dividing by zero is not mathematically allowed, all treatment values were placed in the numerator and divided by control values to calculate a ratio. This ratio was used for all subsequent GWAS calculations. We therefore catalogued GLV10 effects across 171 HapMap lines, quantifying these effects numerically for measuring variation across 14 traits (**Supplementary Table 1**).

The GLV10 peptide had a significant impact across different HapMap lines with some traits responding more strongly than the others (**Figure 1B**). A principal component analysis showed that differences in lateral root traits and nodule traits explained 50% of the variation caused by GLV10p application across all HapMap Lines tested **(Supplementary Figure S1)**. Dependent traits such as nodule number and nodule density clustered together, as expected, while diverging from lateral root traits. Of note were traits 4 (number of elongated LRs) and trait 5 (Total LR number) and their derivatives, Trait 8 (Density of Elongated LRs) and Trait 9 (Density of Total LRs), and less prominently Traits 2 (LR primordia) and Trait 6 (LR primordia density) which contributed to 30% of the observed variation. The nodulation traits clustered slightly separately from the LR traits, with Trait 12 (Nodule number), Trait 15 (Nodule density) and Trait 18 (Nodule zone) contributing most to the observed variation. As seen in Wild Type *M. truncatula* Jemalong A17 plants, a general trend for all root growth traits was a reduction upon GLV10 peptide application. Since GLVs had a repressive effect on all measured traits except root length, the ratios calculated for GWAS had a high frequency of zero values. Consequently, all traits were not normally distributed and skewed to the right upon GLV10 application (**Figure 1D**).

A Pearson correlation test confirmed anticipated relationships between the measured traits, such as an inverse correlation between root length and organ density traits, as well as between nodule position and the subsequent nodule zone length upon GLV10 application (**Supplementary Figure 1**). Additionally, as would be expected, a positive correlation was noted between the number of organs and their respective density measurements. Interestingly, upon peptide treatment, a negative correlation emerged between all nodulation and lateral root traits, a relationship absent under untreated conditions (**Supplementary Figure 1**). This supports the hypothesis that there exists a competition between nodules and lateral roots for space on the primary root and a possible role for GLV10 in resource allocation to either organ type (Bensmihen, 2015).

### Associating genotypic variation with Phenotypic differences across *M. truncatula* Accessions in response to GLV10 peptide

The GLV10 peptide notably influences various root architecture traits in *M. truncatula* (Supplementary Figure Y) (Roy et al., 2024). Of the 14 GWAS trait datasets we evaluated, six revealed significant SNPs and one trait displayed a single significant association when applying a strict qValue, a statistical measure used to account for the false discovery rate (FDR), at a threshold of 0.05 and below (**Supplementary Table 2**). To characterize phenotypic variation, we included the reference line Jemalong A17 in all our experiments for comparison with other HapMap lines. In the Jemalong A17 ecotype, nodule number and density reduced upon GLV10p application (**Figure 2**). While several lines mimicked this reduction, some were unresponsive to the peptide, and a few displayed increased nodule numbers upon peptide treatment (**Figure 2 A, B**). GLV10p application leads to a contraction in the nodulation zone and, when two nodules formed successively, spacing between them reduces (Roy et al., 2024). Many plants in the GWAS population displayed a reduced nodulation zone, but some expanded it (**Figure 2C**). This contraction corresponds with the altered positioning of the initial nodule formed relative to the primary root length. This trait showed variation, with many lines mirroring the effects of A17, while others either remained unaffected or were shifted towards the shoot base (**Figure 2D**). For lateral root characteristics, we scored the number of primordia, emerging lateral roots, and elongated lateral roots. GLV10 peptide application led to a decline in elongated lateral roots and total lateral root density (**Figure 2 E, F**). Few lines increased their lateral root count, while some remained unaffected by the peptide. The data revealed a diverse spectrum of phenotypes within the population, providing an ideal foundation for conducting further statistical analyses using GWAS.

**Figure 2.**
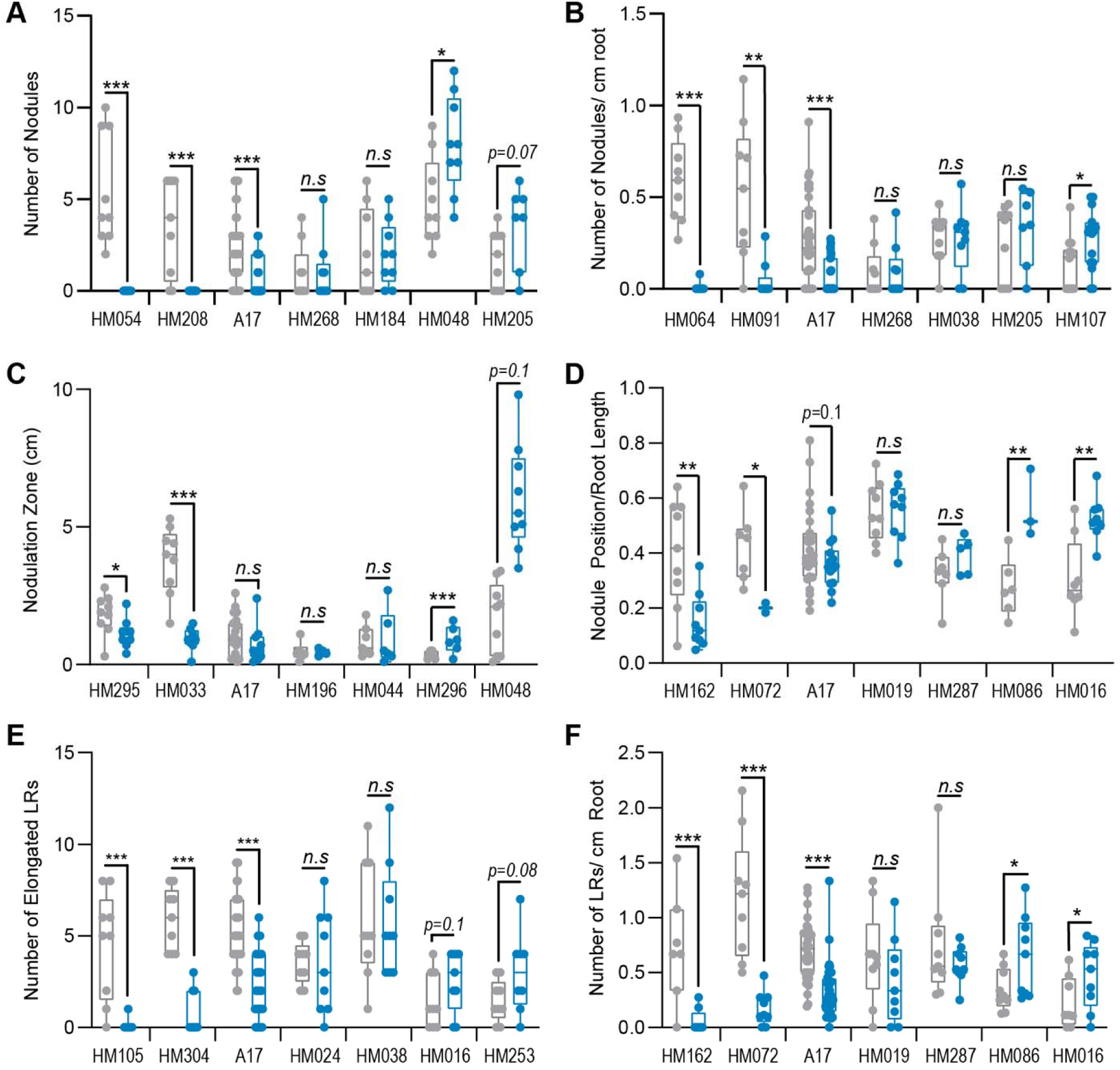
Differential response to GLV10 peptide application across HapMap lines. Boxplots comparing *M. truncatula* hapmap lines treated with and without peptides for A. Nodule Count B. Nodule Density (nodules/cm of primary root) C. Nodulation Zone D. Position of First Nodule Relative to Root Length E. Count of Elongated Lateral Roots F. Lateral Root Density (roots/cm of primary root). Data presented represent lines with phenotypes at the extreme ends of each spectrum. Asterisks represent *p<0.05, \*\**p*<0.01, \*\*\**p*<0.001 using a Student’s t-test. n.s - not significant.

### GWAS revealed a total of 74 unique associations with seven phenotypic traits

GWAS was performed on fourteen traits, using a General Linear Model and seven traits had a total of 82 significant SNPs using a False Discovery Rate (FDR) cutoff q-value of 0.05 and below (**Supplementary Table 2**). Nodule number and nodule density shared eight markers in common, bringing the total to 74 unique SNPs (**Supplementary Table 2**). The quantile-quantile (QQ) plots corresponding to seven traits showed a clear deviation of observed *p*-values from the expected null hypothesis (**Figure 3A**). Nodule density (22 SNPs), nodule number (20 SNPs), and number of elongated LRs (19 SNPs), had the highest number of significant SNPs, followed by relative nodule position 2 (10), total LR density (6), nod zone (3) and LR primordia number (1 SNP) (**Figure 3C, Supplementary Table 1**). The SNPs were distributed across all eight chromosomes and there appeared to be no hotspots for the SNPs to appear (**Figure 3C**). Four SNPs associated with the Nodule Density trait were placed on scaffold regions not yet placed on a chromosome region due to assembly errors, about 62% (47 SNPs) of the SNPs were intergenic, as is expected from several other GWAS studies, 18% (13 SNPs) were present in exons, 10% (7 SNPs) were within introns and 10% (7 SNPs) were within putative regulatory regions i.e the upstream promoter or downstream UTR regions according to MtV5.0 gene models (**Table 1**).

**Figure 3.**
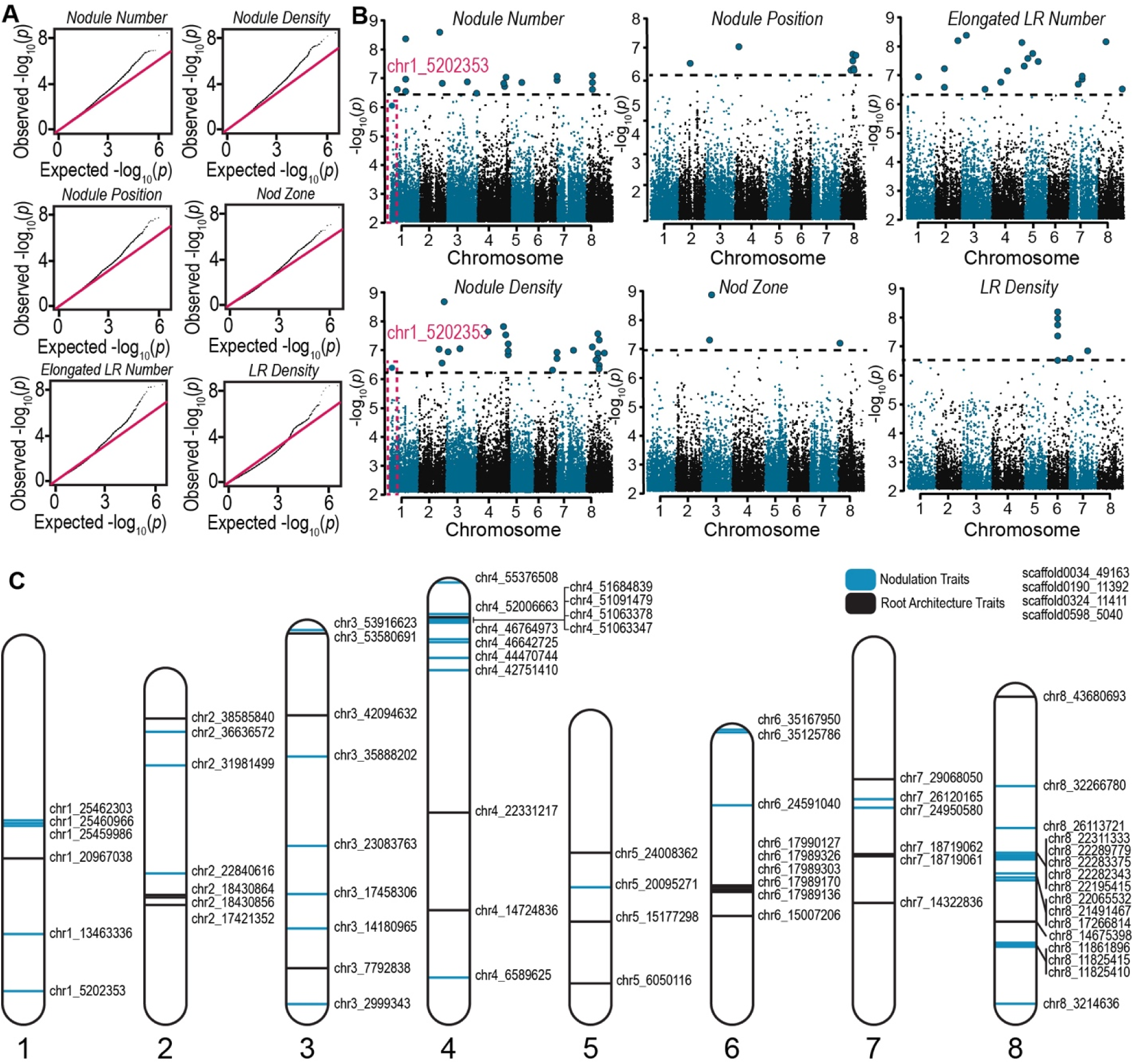
GLV10 peptide treated GWAS study and chromosomal trait distribution. **A.** Quantile-quantile plots showing divergence from prediction for six traits that yielded significant SNPs. **B.** Manhattan plots displaying SNPs associated with the nodule density trait and the highlighted marker chr1_5202353 significant for nodule density. C. Schematic illustration showing positon of 73 SNPs identified using GWAS in *M. truncatula* which were associated with GOLVEN10 control of nodulation and root architecture. Markers in blue are nodulation trait associated while those in black are associated with root architecture. Chromosome size is scaled to data available on Medicago.toulouse.inra.fr.

**Table 1:**
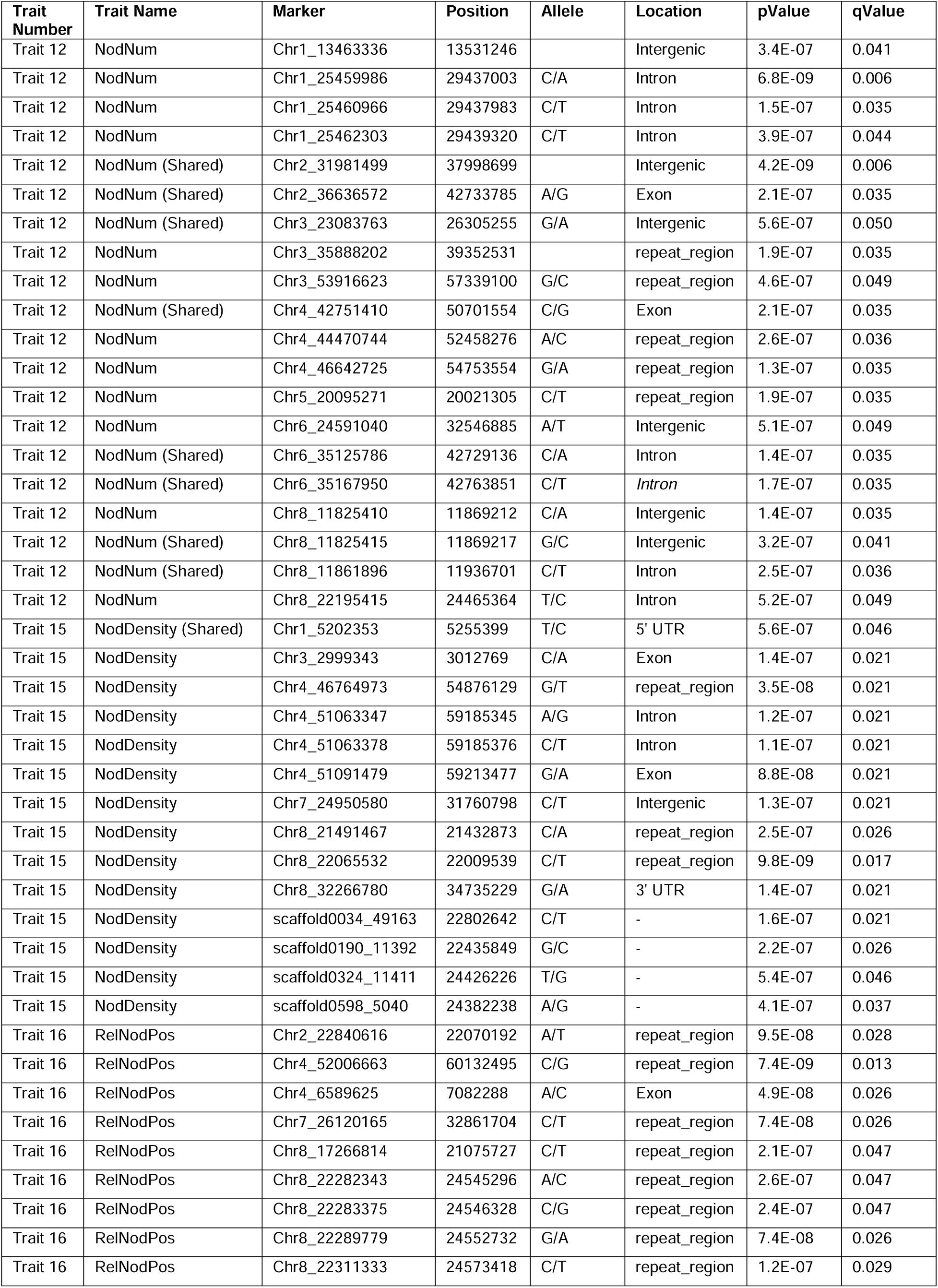
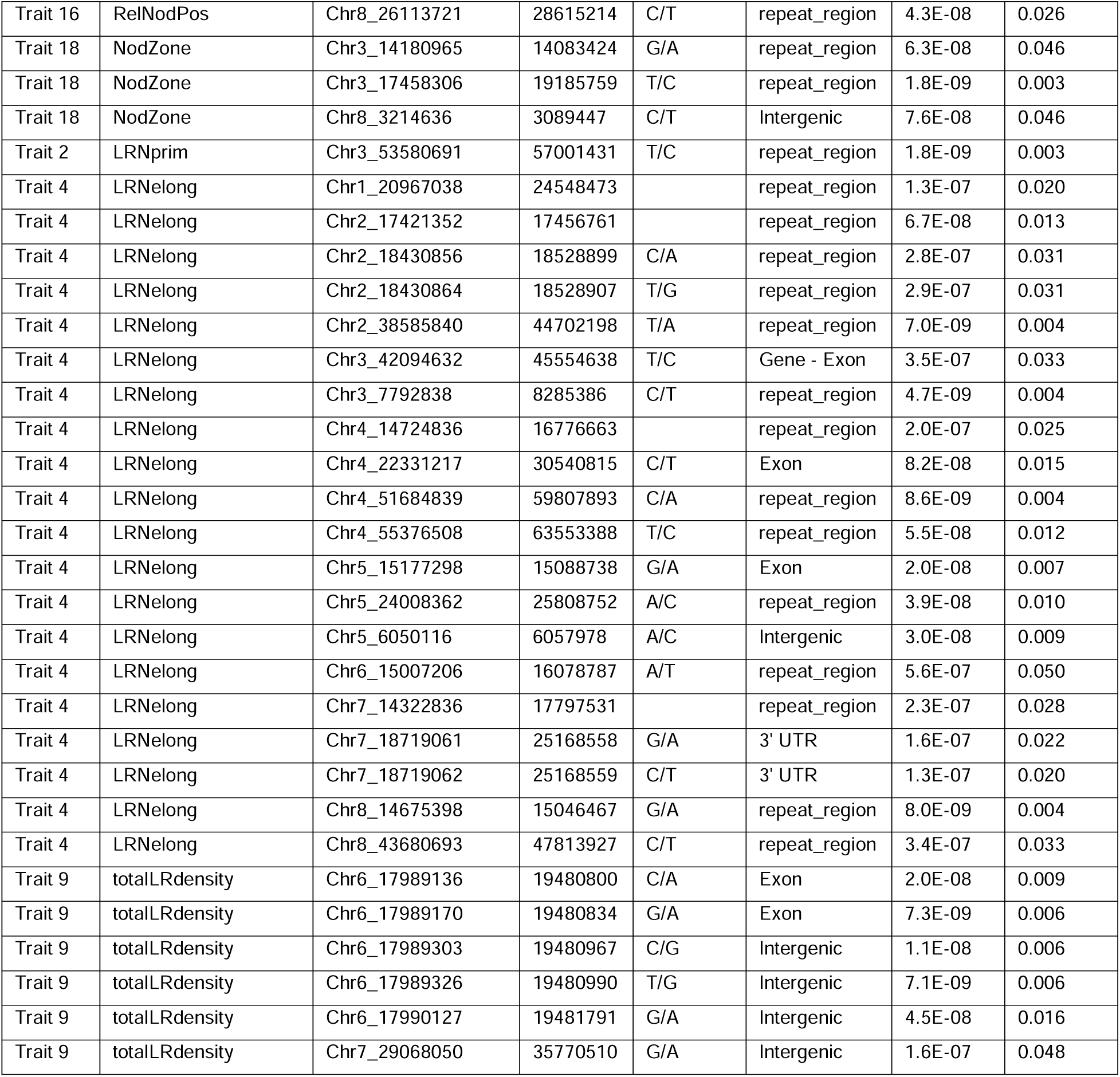
List of SNPs significantly associated with nodulation and root architecture traits investigated in this study. Please see Supplementary Table two for detailed information regarding the SNPs.

| Trait Number | Trait Name | Marker | Position | Allele | Location | pValue | qValue |
| --- | --- | --- | --- | --- | --- | --- | --- |
| Trait 12 | NodNum | Chr1_13463336 | 13531246 |  | Intergenic | 3.4E-07 | 0.041 |
| Trait 12 | NodNum | Chr1_25459986 | 29437003 | C/A | Intron | 6.8E-09 | 0.006 |
| Trait 12 | NodNum | Chr1_25460966 | 29437983 | C/T | Intron | 1.5E-07 | 0.035 |
| Trait 12 | NodNum | Chr1_25462303 | 29439320 | C/T | Intron | 3.9E-07 | 0.044 |
| Trait 12 | NodNum (Shared) | Chr2_31981499 | 37998699 |  | Intergenic | 4.2E-09 | 0.006 |
| Trait 12 | NodNum (Shared) | Chr2_36636572 | 42733785 | A/G | Exon | 2.1E-07 | 0.035 |
| Trait 12 | NodNum (Shared) | Chr3_23083763 | 26305255 | G/A | Intergenic | 5.6E-07 | 0.050 |
| Trait 12 | NodNum | Chr3_35888202 | 39352531 |  | repeat_region | 1.9E-07 | 0.035 |
| Trait 12 | NodNum | Chr3_53916623 | 57339100 | G/C | repeat_region | 4.6E-07 | 0.049 |
| Trait 12 | NodNum (Shared) | Chr4_42751410 | 50701554 | C/G | Exon | 2.1E-07 | 0.035 |
| Trait 12 | NodNum | Chr4_44470744 | 52458276 | A/C | repeat_region | 2.6E-07 | 0.036 |
| Trait 12 | NodNum | Chr4_46642725 | 54753554 | G/A | repeat_region | 1.3E-07 | 0.035 |
| Trait 12 | NodNum | Chr5_20095271 | 20021305 | C/T | repeat_region | 1.9E-07 | 0.035 |
| Trait 12 | NodNum | Chr6_24591040 | 32546885 | A/T | Intergenic | 5.1E-07 | 0.049 |
| Trait 12 | NodNum (Shared) | Chr6_35125786 | 42729136 | C/A | Intron | 1.4E-07 | 0.035 |
| Trait 12 | NodNum (Shared) | Chr6_35167950 | 42763851 | C/T | Intron | 1.7E-07 | 0.035 |
| Trait 12 | NodNum | Chr8_11825410 | 11869212 | C/A | Intergenic | 1.4E-07 | 0.035 |
| Trait 12 | NodNum (Shared) | Chr8_11825415 | 11869217 | G/C | Intergenic | 3.2E-07 | 0.041 |
| Trait 12 | NodNum (Shared) | Chr8_11861896 | 11936701 | C/T | Intron | 2.5E-07 | 0.036 |
| Trait 12 | NodNum | Chr8_22195415 | 24465364 | T/C | Intron | 5.2E-07 | 0.049 |
| Trait 15 | NodDensity (Shared) | Chr1_5202353 | 5255399 | T/C | 5' UTR | 5.6E-07 | 0.046 |
| Trait 15 | NodDensity | Chr3_2999343 | 3012769 | C/A | Exon | 1.4E-07 | 0.021 |
| Trait 15 | NodDensity | Chr4_46764973 | 54876129 | G/T | repeat_region | 3.5E-08 | 0.021 |
| Trait 15 | NodDensity | Chr4_51063347 | 59185345 | A/G | Intron | 1.2E-07 | 0.021 |
| Trait 15 | NodDensity | Chr4_51063378 | 59185376 | C/T | Intron | 1.1E-07 | 0.021 |
| Trait 15 | NodDensity | Chr4_51091479 | 59213477 | G/A | Exon | 8.8E-08 | 0.021 |
| Trait 15 | NodDensity | Chr7_24950580 | 31760798 | C/T | Intergenic | 1.3E-07 | 0.021 |
| Trait 15 | NodDensity | Chr8_21491467 | 21432873 | C/A | repeat_region | 2.5E-07 | 0.026 |
| Trait 15 | NodDensity | Chr8_22065532 | 22009539 | C/T | repeat_region | 9.8E-09 | 0.017 |
| Trait 15 | NodDensity | Chr8_32266780 | 34735229 | G/A | 3' UTR | 1.4E-07 | 0.021 |
| Trait 15 | NodDensity | scaffold0034_49163 | 22802642 | C/T | - | 1.6E-07 | 0.021 |
| Trait 15 | NodDensity | scaffold0190_11392 | 22435849 | G/C | - | 2.2E-07 | 0.026 |
| Trait 15 | NodDensity | scaffold0324_11411 | 24426226 | T/G | - | 5.4E-07 | 0.046 |
| Trait 15 | NodDensity | scaffold0598_5040 | 24382238 | A/G | - | 4.1E-07 | 0.037 |
| Trait 16 | RelNodPos | Chr2_22840616 | 22070192 | A/T | repeat_region | 9.5E-08 | 0.028 |
| Trait 16 | RelNodPos | Chr4_52006663 | 60132495 | C/G | repeat_region | 7.4E-09 | 0.013 |
| Trait 16 | RelNodPos | Chr4_6589625 | 7082288 | A/C | Exon | 4.9E-08 | 0.026 |
| Trait 16 | RelNodPos | Chr7_26120165 | 32861704 | C/T | repeat_region | 7.4E-08 | 0.026 |
| Trait 16 | RelNodPos | Chr8_17266814 | 21075727 | C/T | repeat_region | 2.1E-07 | 0.047 |
| Trait 16 | RelNodPos | Chr8_22282343 | 24545296 | A/C | repeat_region | 2.6E-07 | 0.047 |
| Trait 16 | RelNodPos | Chr8_22283375 | 24546328 | C/G | repeat_region | 2.4E-07 | 0.047 |
| Trait 16 | RelNodPos | Chr8_22289779 | 24552732 | G/A | repeat_region | 7.4E-08 | 0.026 |
| Trait 16 | RelNodPos | Chr8_22311333 | 24573418 | C/T | repeat_region | 1.2E-07 | 0.029 |
| Trait 16 | RelNodPos | Chr8_26113721 | 28615214 | C/T | repeat_region | 4.3E-08 | 0.026 |
| Trait 18 | NodZone | Chr3_14180965 | 14083424 | G/A | repeat_region | 6.3E-08 | 0.046 |
| Trait 18 | NodZone | Chr3_17458306 | 19185759 | T/C | repeat_region | 1.8E-09 | 0.003 |
| Trait 18 | NodZone | Chr8_3214636 | 3089447 | C/T | Intergenic | 7.6E-08 | 0.046 |
| Trait 2 | LRNprim | Chr3_53580691 | 57001431 | T/C | repeat_region | 1.8E-09 | 0.003 |
| Trait 4 | LRNelong | Chr1_20967038 | 24548473 |  | repeat_region | 1.3E-07 | 0.020 |
| Trait 4 | LRNelong | Chr2_17421352 | 17456761 |  | repeat_region | 6.7E-08 | 0.013 |
| Trait 4 | LRNelong | Chr2_18430856 | 18528899 | C/A | repeat_region | 2.8E-07 | 0.031 |
| Trait 4 | LRNelong | Chr2_18430864 | 18528907 | T/G | repeat_region | 2.9E-07 | 0.031 |
| Trait 4 | LRNelong | Chr2_38585840 | 44702198 | T/A | repeat_region | 7.0E-09 | 0.004 |
| Trait 4 | LRNelong | Chr3_42094632 | 45554638 | T/C | Gene - Exon | 3.5E-07 | 0.033 |
| Trait 4 | LRNelong | Chr3_7792838 | 8285386 | C/T | repeat_region | 4.7E-09 | 0.004 |
| Trait 4 | LRNelong | Chr4_14724836 | 16776663 |  | repeat_region | 2.0E-07 | 0.025 |
| Trait 4 | LRNelong | Chr4_22331217 | 30540815 | C/T | Exon | 8.2E-08 | 0.015 |
| Trait 4 | LRNelong | Chr4_51684839 | 59807893 | C/A | repeat_region | 8.6E-09 | 0.004 |
| Trait 4 | LRNelong | Chr4_55376508 | 63553388 | T/C | repeat_region | 5.5E-08 | 0.012 |
| Trait 4 | LRNelong | Chr5_15177298 | 15088738 | G/A | Exon | 2.0E-08 | 0.007 |
| Trait 4 | LRNelong | Chr5_24008362 | 25808752 | A/C | repeat_region | 3.9E-08 | 0.010 |
| Trait 4 | LRNelong | Chr5_6050116 | 6057978 | A/C | Intergenic | 3.0E-08 | 0.009 |
| Trait 4 | LRNelong | Chr6_15007206 | 16078787 | A/T | repeat_region | 5.6E-07 | 0.050 |
| Trait 4 | LRNelong | Chr7_14322836 | 17797531 |  | repeat_region | 2.3E-07 | 0.028 |
| Trait 4 | LRNelong | Chr7_18719061 | 25168558 | G/A | 3' UTR | 1.6E-07 | 0.022 |
| Trait 4 | LRNelong | Chr7_18719062 | 25168559 | C/T | 3' UTR | 1.3E-07 | 0.020 |
| Trait 4 | LRNelong | Chr8_14675398 | 15046467 | G/A | repeat_region | 8.0E-09 | 0.004 |
| Trait 4 | LRNelong | Chr8_43680693 | 47813927 | C/T | repeat_region | 3.4E-07 | 0.033 |
| Trait 9 | totalLRdensity | Chr6_17989136 | 19480800 | C/A | Exon | 2.0E-08 | 0.009 |
| Trait 9 | totalLRdensity | Chr6_17989170 | 19480834 | G/A | Exon | 7.3E-09 | 0.006 |
| Trait 9 | totalLRdensity | Chr6_17989303 | 19480967 | C/G | Intergenic | 1.1E-08 | 0.006 |
| Trait 9 | totalLRdensity | Chr6_17989326 | 19480990 | T/G | Intergenic | 7.1E-09 | 0.006 |
| Trait 9 | totalLRdensity | Chr6_17990127 | 19481791 | G/A | Intergenic | 4.5E-08 | 0.016 |
| Trait 9 | totalLRdensity | Chr7_29068050 | 35770510 | G/A | Intergenic | 1.6E-07 | 0.048 |

### Gene Ontology Analysis of significant associations and candidate gene selection

To identify potential causative genes linked to the SNPs of interest, we scanned a 100 Kb window of the genome which included 50 kb on either side of each SNP. This resulted in a total of 973 genes with 727 putative protein coding genes and 246 unannotated open reading frames using both *M. truncatula* V5 and V4 genome annotations. A Gene Ontology (GO) enrichment analysis using all 973 genes found a significant enrichment of ‘peptidase activity’ using the molecular function scope. Further, we grouped all GO categories with similar functions into 12 categories using ‘CateGOrizer’ and manual curation (Na et al., 2014). This analysis revealed that the most overrepresented genes were categorized as enzymes, with Transferases (24) and Hydrolases (22) being the most prevalent, followed by Endopeptidases and Kinases/Phosphatases **(Figure Z)**. Since we were particularly interested in signaling genes, we individually categorized all 973 genes into four categories – Signals (23) included peptides both post-translationally modified peptides and cysteine-rich peptides, Receptors (2), Signal Transducers (52) included the kinases and phosphatases involved in signal relay, and, Regulators (37) which included most of the transcription factors based on their V4 or V5 annotations. Since the broad 100 Kb window included several genes that are simply adjacent to the causative gene of interest, we next looked at whether the expression of any of these genes was also regulated by the peptide GLV10 using published RNA seq data. This helped narrow down the list to 23 genes that were transcriptionally also regulated by application of the peptide GLV10. Of the two receptors, only 1 gene encoding the MtDWARF14 strigolactone receptor was also significantly regulated by the hormones. This gene encodes for a non-canonical a/b hydrolase gene that perceives the strigolactone.

Known nodulation genes included (associated traits)

This SNP was associated with the nodule density traits at the q = 0.05 and below level 0.15 for the nodule number trait. Intriguingly, the first category included the receptor of the sesquiterpene Strigolactone which acts as a phytohormone regulating various root growth traits. The causal SNP on chromosome 1 associated with nodule density (chr1_5202353, q-value 0.046) and nodule number (q-value 0.151) was 102 bp upstream of the coding region of the gene for strigolactone receptor *MtDWARF14 (D14)* (MtrunA17_Chr1g0152551, Medtr1g018320), two bp upstream of a putative 8 bp TATA binding site (TATAAATA) which is a core promoter element involved in transcription initiation (**Figure 4, Figure S1**). Most lines associated with this SNP showed no decrease in nodule density upon GLV10 peptide treatment (**Figure 4**). This may be because variations in the TATA box can affect binding of RNA Polymerase II to the core promoter region that drives not only basal gene expression but also environment-dependent gene expression (Grace et al., 2004). The TATA box, identified by its consensus sequence of eight base pairs, “TATA(A/T)A(A/T)(A/G),” typically resides between 30 and 70 base pairs upstream from the transcription start site in plant promoters (Brooks et al., 2023) but some studies identified the 13 bp TCACTATATATAG as the consensus sequence in promoters of highly expressed plant genes (Kiran et al., 2006). These GC-rich TATA extension sequences that flank the TATA motif are thought to govern the formation of alternative transcriptional complexes needed in different physiological settings and alter the rate and stability of TATA binding protein and TFIIB transcription factor binding to the TATA box to recruit RNA Polymerase II (Wolner and Gralla, 2001; Kiran et al., 2006; Shi and Zhou, 2006). In transient b-Glucuronidase expression assays carried out in *Nicotiana benthamiana*, a C_2_ – T mutation, i.e. a mutation of the second residue in the 13 bp consensus sequence, as is the case in the chr1_5202353 SNP, decreased transcription substantially (Kiran et al., 2006). To eliminate the possibility that the chr1_5202353 SNP appears in plants that simply make more nodules, we also ran a GWAS for the nodule density trait in the untreated HapMap lines alone and this SNP was absent (**Supplementary Table 2**). This confirmed that the chr1_5202353 SNP is associated with sensitivity to the GLV10 peptide and not nodule density alone. Two accessions with the chr1_5202353 SNP still retain sensitivity to the GLV10 peptide (HM166 and HM198). This could be because there exist other genetic factors in the genomic backgrounds of these two lines that are required for nodule formation which is a complex trait. Interaction between the strigolactone pathway and other hormone signals determine the final outcome.

**Figure 4:**
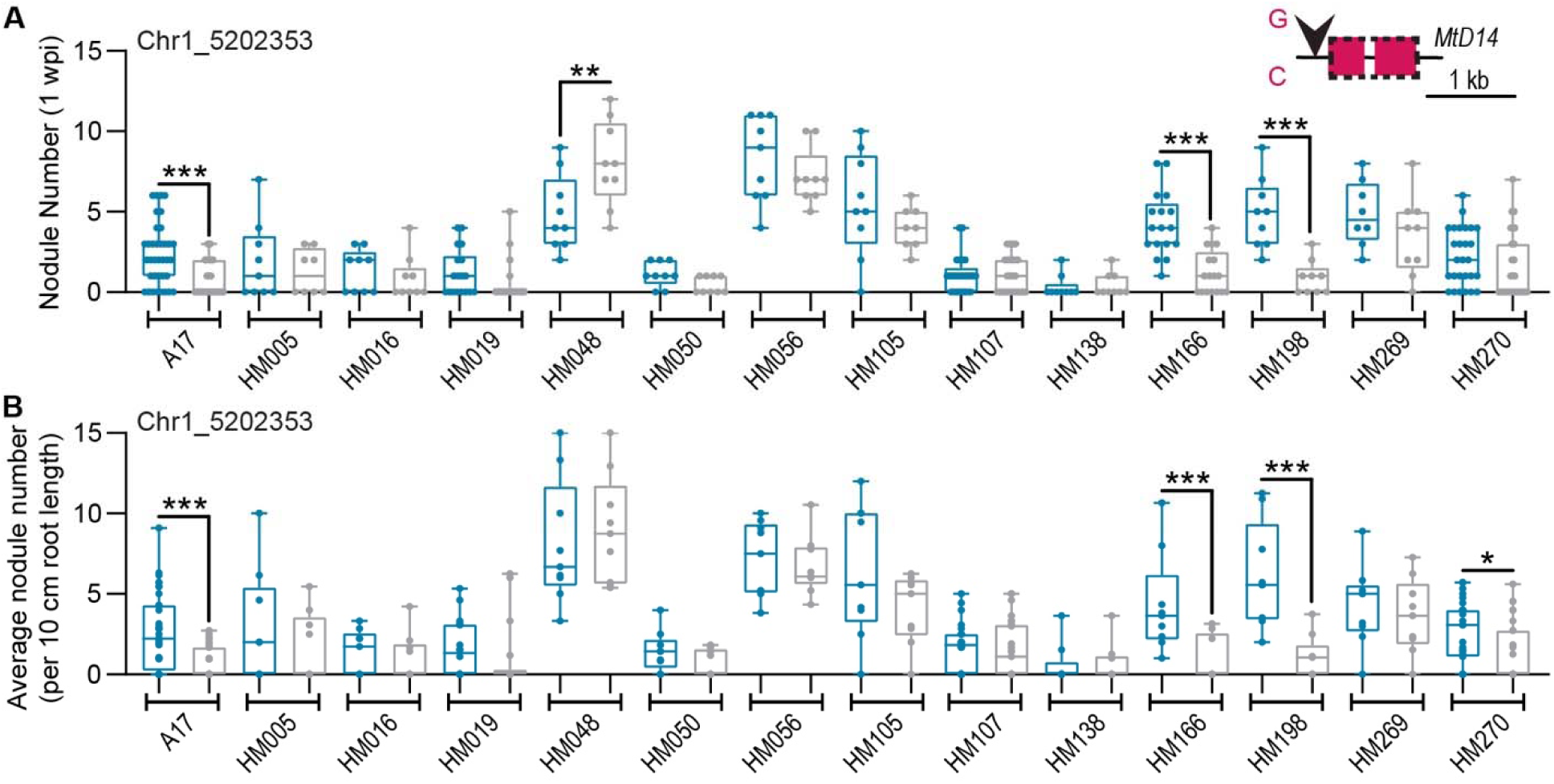
Nodulation phenotypes of thirteen *M. truncatula* HapMap lines associated with a SNP on chromosome 1. Response of HapMap lines to GLV10p in lines associated with the A. nodule number trait and B. Nodule density trait associated with marker chr1_5202353. Asterisks indicate \*\**p*<0.01 using an ANOVA-protected Sidak’s multiple comparison test. Inset shows position of SNP in the upstream promoter region of the *MtDWARF14* gene.

### GLVs differentially regulate strigolactone biosynthesis, transport and signaling

In parallel with GWAS, we investigated Medicago transcriptomic responses to GLV10p treatment, using quantitative Reverse Transcriptase PCR (qRT PCR). Tthe q-PCR data provided an independent line of evidence that corroborated GWAS results and indicated that the GLV signaling pathway regulates key strigolactone (SL) biosynthesis, transport and signaling genes. SLs are both perceived and hydrolyzed by the α/β hydrolase enzyme DWARF14 (D14) (REF). Binding of SL leads to a conformational change in the D14 complex with the F-Box protein MORE-AXILLARY 2 (MAX2), which degrades SUPRESSOR OF MORE AXILLARY LIKE (SMAX-LIKE) repressor proteins, allowing expression of SL-responsive genes (Mashiguchi et al., 2021). Homologs of all three SL-signaling genes, *MtD14*, *MtMAX2* and *MtSMXL6* were up-regulated by GLV10p application in *M. truncatula* (**Figure 5**), while expression of SL biosynthesis genes (*MtD27a* and *MtCCD8*) was repressed (**Figure 5**), consistent with previous data suggesting SL biosynthesis is under a negative feedback loop regulated by its own signaling (Chevalier et al., 2014). Additionally, the transporter responsible for root acropetal transport and exudation of SLs, *MtABCG59*, was strongly repressed by GLV10p treatment (**Figure 5**) (Banasiak et al., 2020). SL signaling acts synergistically with karrikin signaling on many developmental processes, including primary root growth and LR density, and we found that *MtKAI2b* and *MtSMAX1* were induced by GLV10p (**Supplementary Figure S2**) (Villaécija-Aguilar et al., 2019). Karrikin is a smoke derived signal, the *in-planta* counterpart of which is currently unknown (Swarbreck et al., 2020). Moreover, overexpression of *MtGLV10 in* chimeric hairy roots also induced SL signaling genes in hairy roots of *M. truncatula* 21 dpi with *S. meliloti* Rm1021 in line with peptide application studies (**Figure 5B**). This pointed to an interaction between strigolactone and GLV signaling pathways *in planta*.

**Figure 5:**
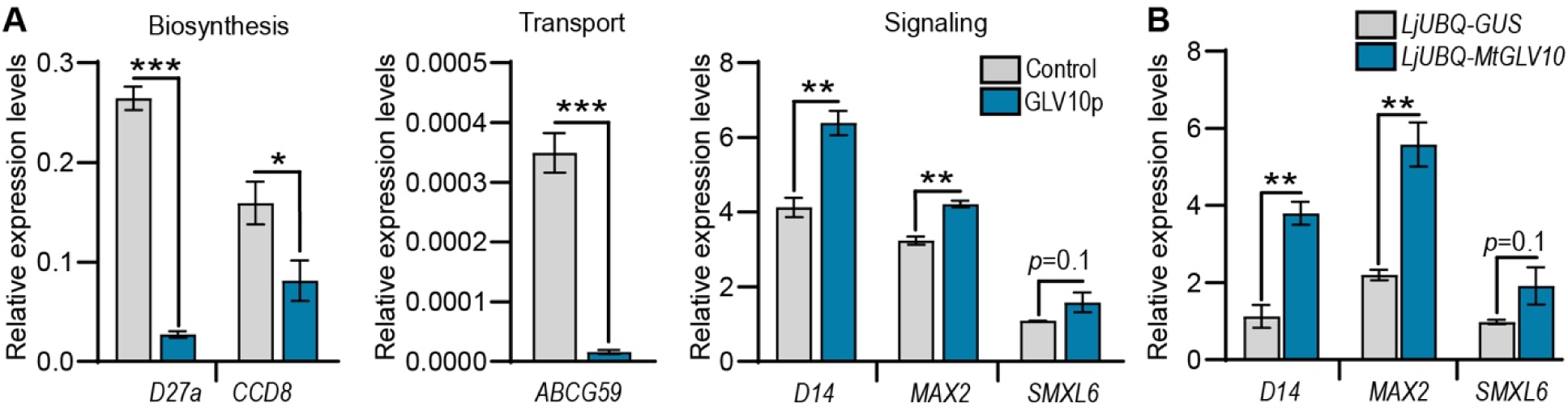
Application of GOLVEN10 peptide or overexpression of GOLVEN10 prepropeptide alters expression of strigolactone pathway genes. **A** Strigolactone signaling, transport and biosynthesis gene expression levels as measured by quantitative RT-PCR following GLV10p peptide application. Error bars depict SEM. Student’s *t*-test \**p*<0.05, \*\**p*<0.01, \*\*\**p*<0.001. **B.** qRT-PCR determination of strigolactone signaling genes in *MtGLV10* overexpression hairy root lines of *M. truncatula* Jemalong A17 lines 21 dpi with Rm1021. Error bars represent SEM. Each treatment comprises of three biological replicates. Asterisks represent \**p*<0.05, \*\**p*<0.01 using a Student’s *t*-test. All qRT-PCR data displayed were normalized using two housekeeping genes *UBIQUITIN* and *PTB*.

### Functional characterization of DWARF14 overexpression and mutant lines

To understand the within-nodule spatial expression pattern of MtD14 we cloned a 3 Kb region upstream of the translation start site. The candidate gene associated with chromosome 1 marker encoding *MtD14* showed an expression pattern overlapping with that of MtGLV10 and was expressed at nodule initiation sites (**Figure 6**), later becoming meristematic as well as in root tips and lateral roots (**Supplementary Fig. S7**). Since the hapmap lines with a putatively deregulated *D14* gene showed altered response to the GLV10 peptide, we sought to investigate the phenotypic consequences of a total loss or gain of *D14* gene function. Overexpression of *MtD14* in *M. truncatula* hairy roots, suppressed nodule number by 30% upon inoculation with Rm1021, comparable to *GLV* overexpression effects (**Figure 6B**). Further, three independent *Tn1* mutant alleles of *d14-1, d14-2, d14-3* showed a 50% increase in nodule number as well as lateral root number (**Figure 6C, Supplementary Figure S2**). When we normalized the number of nodules to the total root length upon scanning roots, we found that the density of nodules pr cm root was also higher in *d-14* mutants (**Figure 6**). This established that D14 is required for optimal nodule initiation and density in *M. truncatula*.

**Figure 6:**
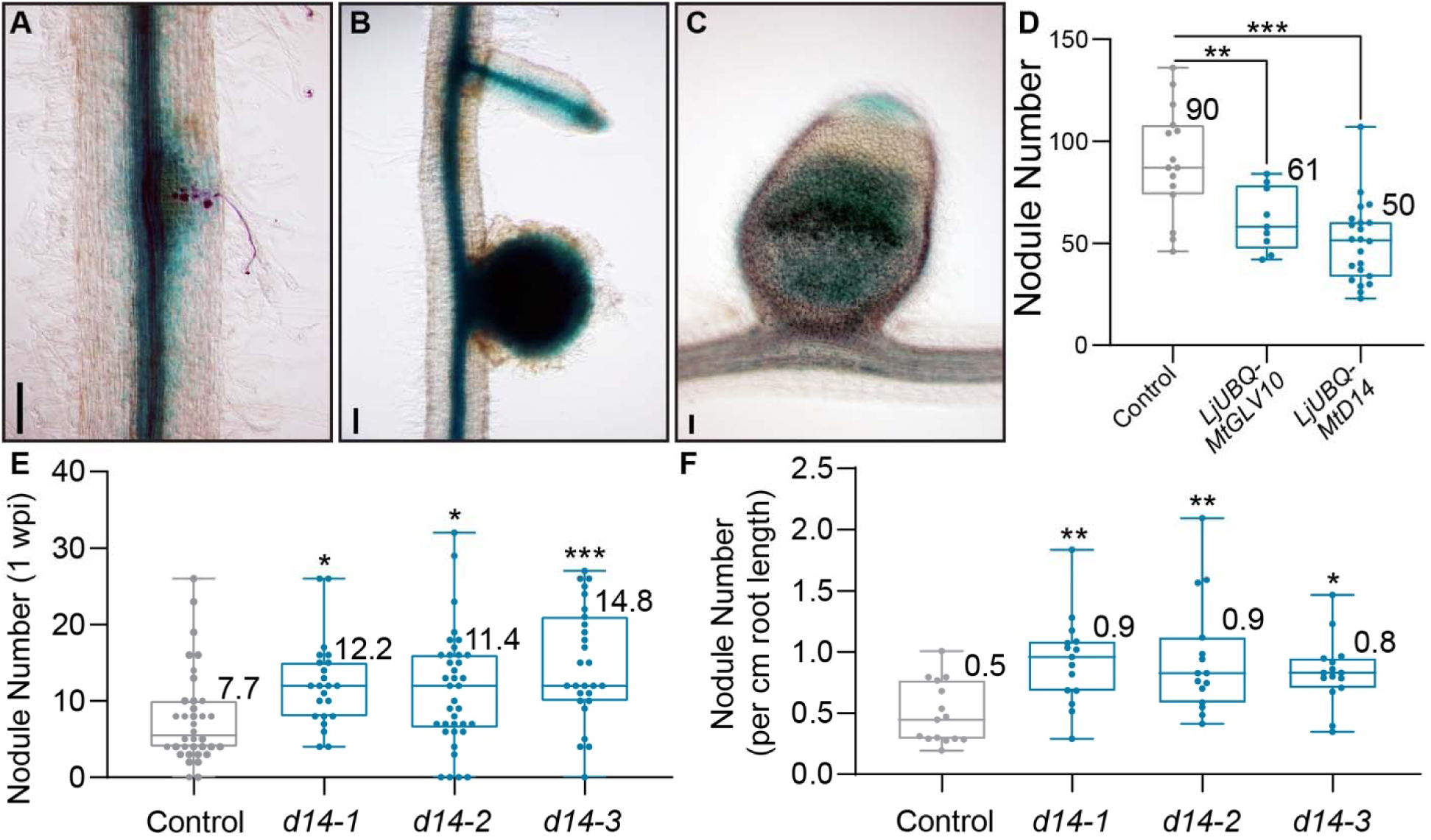
Functional characterization of the MtDWARF14 gene. Spatial expression pattern of 3 kb upstream of the *MtDWARF14* gene in *M. truncatula* hairy roots infected with *S. meliloti* Rm 2011 A. 4 days post inoculation (DPI) B. 10 DPI C. 28 DPI. At least six plants per time point were assessed. D. Overexpression of the *MtGLV10* gene and the *MtDWARF14* gene repress nodulation. E. Boxplot showing nodule numbers on three independent *Tnt1* mutant alleles in *d14* mutants compared to wild type R108 one week post inoculation with *S. meliloti* Rm1021. F. Three independent *Tnt1* mutant alleles in *d14* form more nodules per cm root compared to wild type R108. All experimental groups had at least 10 individual plants. Statistical significance was calculated using an ANOVA test followed by a posthoc Dunnett’s multiple comparison test. Asterisks indicate \**p*<0.05, \*\**p*<0.01, \*\*\**p*<0.001.

### *DWARF14* is required for GLV10 induction of two *PLETHORA* transcription factors

Synthetic strigolactones such as GR24 can mimic the *in planta* effects of the hormone when applied exogenously (REF). To compare effects of strigolactone (GR24), and GLV10p we tested various concentrations of these hormones on nodulation (**Figure 7A, B**). In our system, high concentrations of both hormones suppressed nodule formation (**Figure 7 B**); although at 1 nM, GR24 but not GLV10 had a slight stimulatory effect on nodule number, as reported for *Pisum sativum* and *M. sativa* (REF). This could be due to effects of strigolactones on rhizobial infection as their biosynthesis is upregulated in infected root hairs of Medicago (Breakspear et al., 2014).

**Figure 7:**
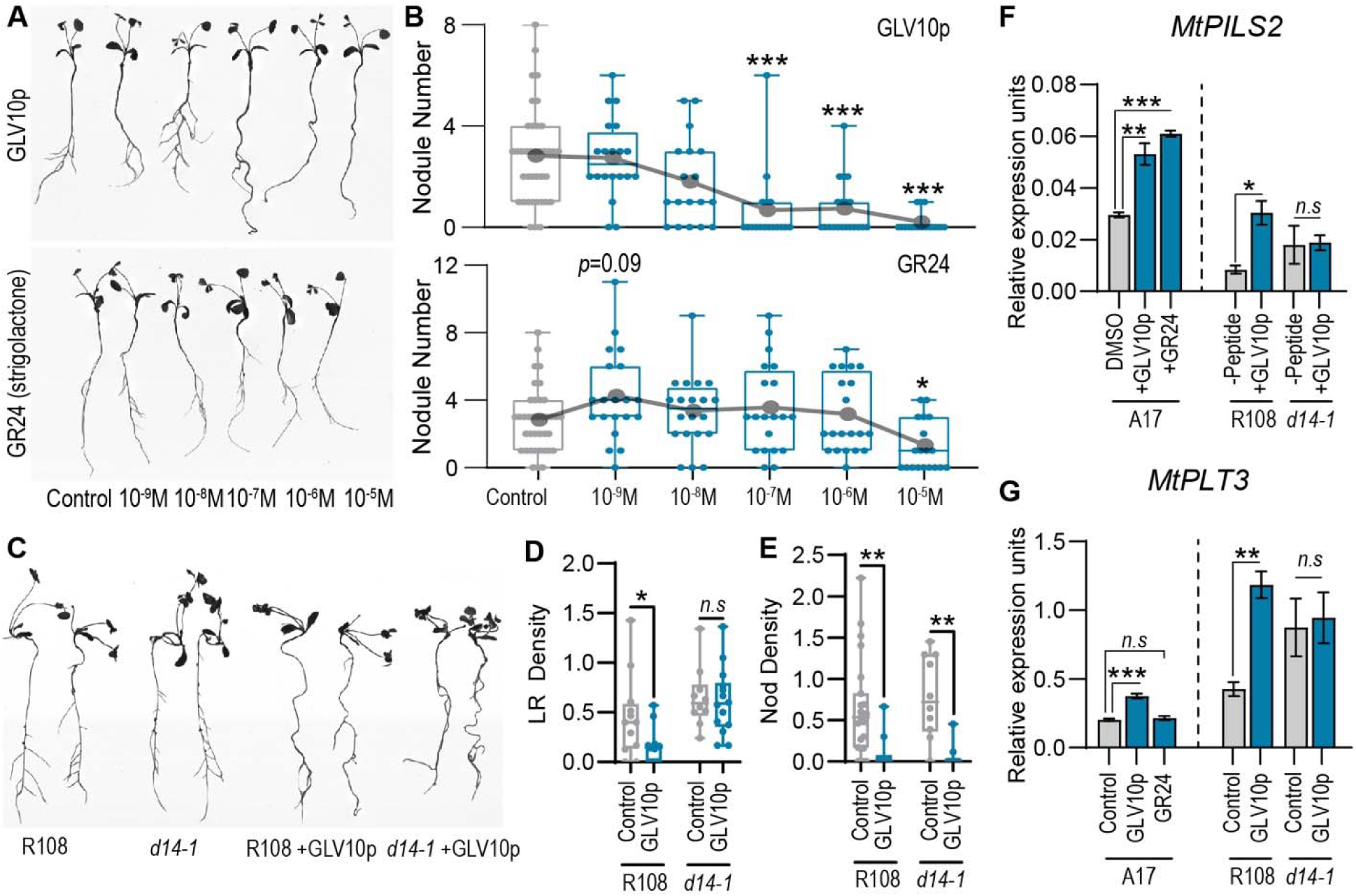
Comparisons between GOLVEN10 and strigolactone signaling pathways. **A** Representative root scans showing seedling morphology 13 days post growth on B&D low-N media containing GLV10p, strigolactone (rac-GR24) at the concentrations indicated. **B.** Number of nodules 10 dpi with Rm2011 dsRed at the concentrations of chemicals indicated. Asterisks indicate \*\**p*<0.01, \*\*\**p*<0.001 using an ANOVA-protected Dunnett’s multiple comparison test. **C.** Representative root scans showing *M. truncatula* morphology 17 days post growth on B&D low-N media containing GLV10p at the concentrations indicated **D.** Boxplot showing number of lateral roots per cm of primary root upon application of GLV10p. Data were combined from two identical experiments. **E.** Boxplot comparing number of nodules per cm of primary root of WT R108 and *d14-1* mutant upon application of GLV10p. Data were combined from two identical experiments. Student’s t-test \**p*<0.01, \*\**p*<0.05, \*\*\**p*<0.001 F**, G** Expression of the two transcription factors *MtPLT3* and *MtPLT5* in three-day old seedlings treated with GLV10p or the synthetic strigolactone rac-GR24 for three hours compared to their respective controls. Expression of the same genes in seedling roots of WT R108 and *d14*-1 mutant lines upon GLV10p treatment for three hours. Each biological replicate comprises of 20 seedlings each and three replicates were used per treatment. Error bars represent SEM. Asterisks represent \**p*<0.05, 690 \*\**p*<0.01, \*\*\**p*<0.001 using a Student’s t-test.

To investigate a potential link between MtGLV10 and strigolactone signaling, we verified that the *Tnt1*-insertion mutant of *D14* in Medicago (*d14-1*) was a null mutant (**Supplementary Figure S1**). *D14* transcript could not be detected in roots of this mutant allele (**Supplementary Figure S1**). Moreover, the homozygous *d14-1* mutants were dwarfed, exhibited a hyperbranching shoot phenotype, and initiated a higher number and density of lateral roots, phenotypes characteristic of *d14* mutants in Arabidopsis (**Supplementary Figure S2**). The *d14-1* null mutant initiated LRs higher on the primary root compared to WT and had a larger zone over which LRs were initiated (**Supplementary Figure S2**). The *d14-1* mutants also initiated 50% more nodules at 7 dpi with Rm1021, which was sustained at 28 dpi in soil (**Fig. 7e**). Since there was no difference in the number of nodules initiated on the primary root alone, we hypothesized that the *d14-1* mutant initiated more nodules due to the higher number of LRs. To test this, we scanned nodulated roots at 7 dpi and measured the total length of each root using RhizoVision and counted nodule numbers (**Supplementary Figure Y**). The number of nodules initiated per cm root (**Supplementary Figure Y**), and the zone over which these nodules were formed was 3x higher in *d14-1* compared to WT R108 (**Figure 7**). These mutant phenotypes were opposite to those invoked by GLV10p application for six traits (i.e., nodule number, nodule position, nod zone, LR density, LR position, LR zone; **Supplementary Figure X**). The *d14-1* mutant was less sensitive to GLV10p than the WT, and often failed to increase root length in response to 50 nM GLV10p (**Supplementary Figure X**). Although there was a high degree of variability, *d14-1* mutants exhibited no change in LR density or LR zone upon GLV10p treatments at 50 nM. While, nodule density and nodule zone were still sensitive to 50 nM GLV10p, 14 dpi with Rm2011. This high variation in GLV10 effect on organ number in the *d14* mutant combined with induction of *D14-like* gene expression by the peptide suggests that higher order mutants may be required. Not only DWARF14 but the Karrikin Signaling pathway involving DWARF14-like genes are also required for formation of lateral organs.

We hypothesized that both strigolactone and GLV10p dissipated auxin gradients, thereby repressing lateral organ priming. Indeed, both GLV10p and GR24 robustly and reproducibly induced the auxin efflux transporter *MtPILS2 (PIN LIKE 2)* gene in three independent experiments (**Figure 7F**). PILs act by effluxing auxin into the endoplasmic reticulum such that less nuclear auxin is available for transcriptional induction of auxin-responsive transcription factors (Sauer and Kleine-Vehn, 2019; Waidmann et al., 2023). GLV10p induction of *MtPILS2* required the *MtD14* receptor (**Figure 7 F**). Since PLETHORA transcription factors are required for nodule initiation and are induced by GLV10p, we tested whether their induction was D14 dependent (Franssen et al., 2015). Indeed, *d14-1* seedling roots treated with GLV10p failed to induce *MtPLT3* and *MtPLT5* (**Figure 7E, Supplementary Figure Z**). These data suggest a model in which the GLVs and SLs interfere with auxin availability by affecting *MtPILS2* and thereby affect transcription of the *MtPLT* transcription factors.

### Medicago Accessions with DWARF14 SNPs correlate with regions that experience a higher frequency of frost days

Gene-environment interactions (GxE) are found to play important roles in plant adaptation to local bio-geochemical variables. The calculated heritability of all traits analyzed in this study were low, registering at under 10%. This can be attributed to the artificial environmental selection pressure exerted by the GLV10 peptide, which was used to uncover genes regulated by the peptide. However, the SNP chr1_5202353 was retained in closely related plants within one clade of a phylogenetic analysis based on kinship of all 171 lines used in this study (**Figure 8**). The SNP on chromosome 1 chr1_5202353 is shared by 13 Medicago lines that cluster within a clade of lines collected from the coastal plains of the Maghreb region of north Africa that includes the Algeria-Morroco-Tunisia region (**Figure 8**) (Yoder et al., 2013). This suggests the SNP might have arisen in a common parental line. The presence of the SNP in two lines from Portugal and Spain may be explained by the fact that Medicago despite being an inbreeding plant does outcross occasionally leading to gene escape (Jullien et al., 2021). We therefore sought to compare our findings with previously conducted GWA studies on 10 climatic variables in regions where the Medicago Hapmap population was collected (Guerrero et al., 2018). Of the 202 accessions reported by Guererro et al., 130 overlapped with lines included within this study. We therefore ran a Pearson correlation analysis of the presence of the SNP with all ten climatic variables. We found that only one trait showed a low but significant correlation with the chr1_5202353 SNP (*r*≥0.2, *p*≤0.05). Medicago accessions that grew in locations which saw a higher frequency of frost days more frequently retained the DWARF14 SNP suggesting it may play a role in providing a survival advantage in these regions.

**Figure 8:**
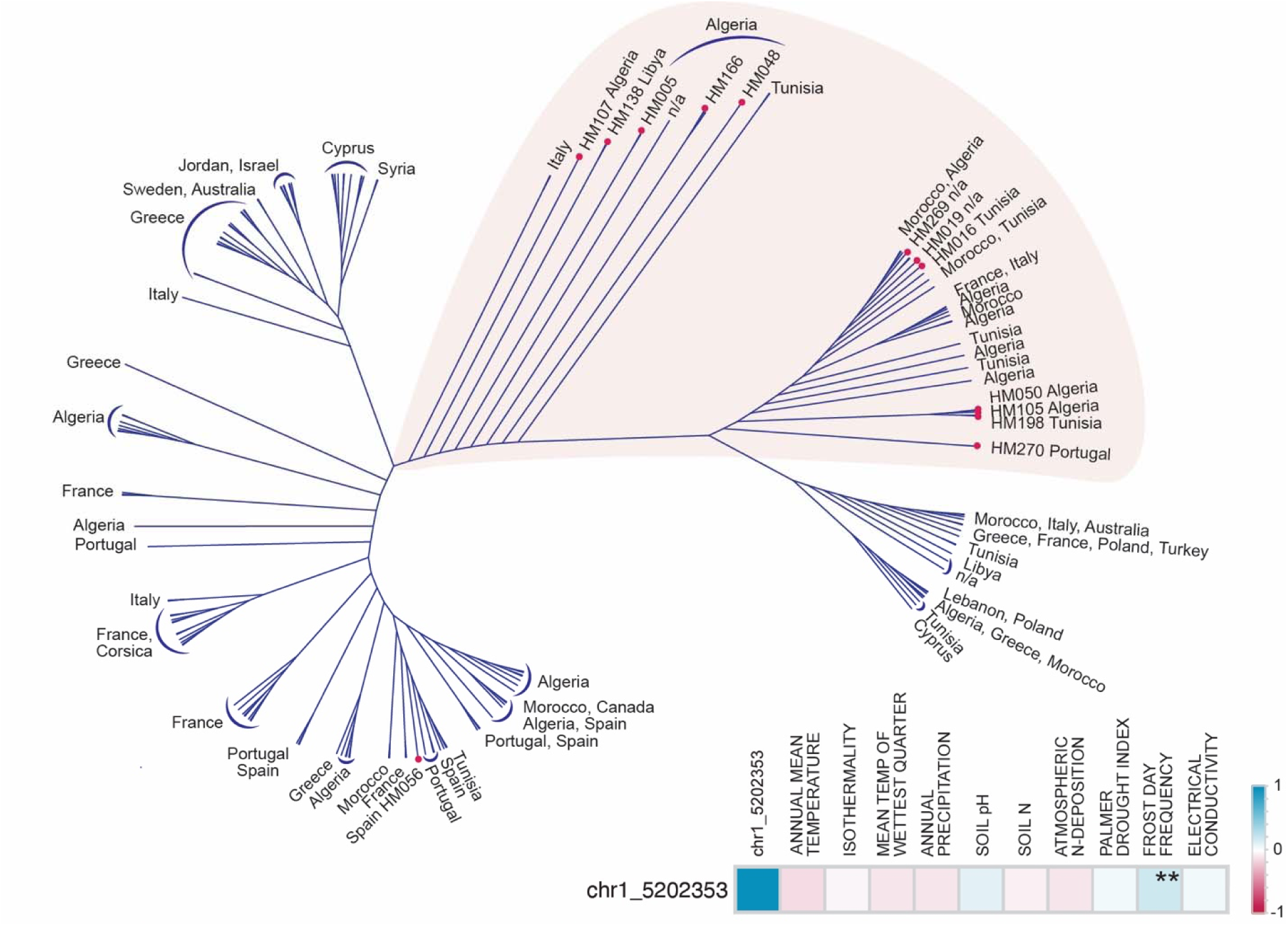
Presence of the *MtDWARF14* associated SNP across *M. truncatula* HapMap lines. Phylogenetic tree showing interrelatedness between all 227 *M. truncatula* HapMap lines generated using complete genomic data in Tassel 5. Blue brackets indicate a group of lines that come from the same region. Magenta dots are shown next to the lines with the candidate SNP on chromosome 1 upstream of the *MtDWARF14* gene.

## DISCUSSION

Root nodule symbiosis enables legume crops to reduce dependence on external fertilizers by facilitating effective nitrogen acquisition in specialized root lateral organs called nodules. At the same time, plants can regulate the proliferation of lateral roots in nitrogen-rich patches as an alternate means to uptake nitrogen. Therefore, studying genetic factors that control these traits is important to be able to engineer plants that are adaptable to different soil types with heterogenous nitrogen availability. The peptide hormone, GLV10, is a critical player in this context. It acts as a powerful growth modulator, controlling both lateral root and nodule formation, particularly under nitrogen-deprived conditions (Roy et al., 2024).

In this study, we developed a thorough compendium of GLV10 peptide responses across 172 *M. truncatula* hapmap accessions. In the majority of the lines, the response closely mimicked that of the reference genome, Jemalong A17 indicating that A17 responses provided a good representative of peptide responses (Supplementary Table X). While RNA sequencing and other estimation tools can provide insight into genes regulated by GOLVEN peptides within one genome, typically the reference genome Jemalong A17, they do not capture the genetic basis that provide adaptive strategies under different environmental conditions. Genome Wide Association Studies are therefore a powerful strategy to capture the genetic variations associated with complex traits and elucidate the underlying molecular mechanisms driving phenotypic diversity. We effectively used this panel in controlled environments to examine the underlying basis of GLV10 peptides on Medicago nodulation and root development, marking a novel approach in the field.

The natural variation in response to the GLV10 facilitated a robust GWAS analysis which yielded 74 SNPs for seven traits. This number is considerably higher than those found for the hormones IAA and Cytokinin using GWA studies in Arabidopsis which yielded five and one significant SNP association respectively under similar controlled conditions (Ristova et al., 2018). We further focused on SNPs associated with signaling related genes and identified the strigolactone receptor DWARF14 as a gene involved in mediated GLV10 responses. Given that *in planta* strigolactones are also required for root architecture and exogenous SLs for interactions with beneficial microbes, we chose to investigate this further during root nodule symbiosis (Akiyama et al., 2005; Foo and Davies, 2011; Muller et al., 2019). Strigolactone is synthesized by sequential action of three enzymes DWARF27 (D27), CAROTENOID CLEAVAGE DIOXYGENASE 7 (CCD7) and CCD8 which forms the SL precursor carlactone (Mashiguchi et al., 2021). The non-canonical receptor DWARF14 is a cytoplasmic alpha/beta fold containing hydrolase that upon hydrolyzing strigolactone molecules binds the F-box protein MAX2 which causes a change in structure and consequently allosteric activation of D14 downstream signaling (Bürger and Chory, 2020). In planta concentration of GOLVENs is very low and therefore maybe d14 mutants still retain sensitivity to this peptide.

Nodule density is an important trait to study because it determines the total number of nodules that can be supported by a plant root even if the root system is not able to expand. Plants that have a higher nodule density can provide advantages to plants for example under waterlogged soil conditions when the root system is stunted. Downstream of the central symbiosis transcription factor NIN, localized auxin maxima formed at nodule primordia are crucial for organogenesis, and GLVs inhibit organ formation by disrupting auxin gradients. They do so by inducing genes encoding auxin efflux transporters, and, deregulating auxin signaling. This effect is strikingly similar to that of strigolactone, which inhibits adventitious root formation, deregulates auxin canalization as mediated by PIN transporters, and suppresses auxin signaling (Koltai, 2015; Jiang et al., 2016; Zhang et al., 2020). GOLVEN peptide application as well in vivo overexpression induced components of the strigolactone signaling pathway (Figure 5). Even though at high concentrations the SL analog GR24 suppressed nodule organogenesis, we observed that at low concentrations (1 nM), SLs exerted a modest stimulatory effect on nodule number (Figure 7). This phenomenon may be linked to rhizobial infection, especially considering the induction of SL biosynthesis genes in infected root hairs (Breakspear et al., 2014). Our work demonstrates that an intact strigolactone signaling pathway is required for GLV induction of *MtPLT3* and *MtPILS2* expression (**Figure 7 F,G**). Orthologs of PLETHORA genes are crucial for nodule formation in Medicago, rhizotaxis and lateral organ positioning in Arabidopsis (Matsuzaki et al., 2010; Hofhuis et al., 2013; Franssen et al., 2015). Considering that SLs are known to regulate phyllotaxis and rhizotaxis, their role in modulating noduletaxis, which is under the control of GOLVENs, aligns with these established functions (Roy et al., 2024). GLV peptides require the SL signaling pathway to alter LR density during early stages of organogenesis (**Figure 7**). Together SL and GLV regulatory networks amplify the signal to inhibit auxin-mediated lateral organ priming, positioning and initiation until the plant has overcome N-deficiency stress (**Figure 7**). Given GLV10s induction of Karrikin signaling receptor, *MtKAI2*, as well (**Supplementary Figure Z**), the absence of an effect at the phenotype level could be due to redundancy of these signaling pathways.

The impact of frost on root systems is poorly studied, yet it is substantially detrimental to global agricultural productivity (Ambroise et al., 2020). Indeed, number of nodules per root weight (surrogate measure for nodule density) reduces with lower soil temperatures and more frequent frost days (Johnson and Rumbaugh, 1986; Junior et al., 2005). This may be due to direct effects on rhizobial growth in cold temperatures, reduced root hair infection and nodule initiation, slower nodule expansion and nitrogen fixation as well as an indirect effect on nodulation due to an overall reduction in plant growth. Frost tolerance refers to a plant’s ability to survive short periods of below freezing temperatures as opposed to cold tolerant plants that can survive extended periods of freezing temperatures. The SNP on chromosome 1 chr1_5202353 is associated with a deregulated DWARF14 gene, which potentially could lead to a greater number of nodules or secondary lateral roots per unit root length (Figure 5). It raises the possibility that this group might exhibit a decreased sensitivity to the GOLVEN-mediated inhibition of nodulation and LR formation, to acquire a survival advantage in these regions. Belowground frost primarily destroys fine roots, which are vital for the uptake of water and minerals (Ambroise et al., 2020). The strigolactone pathway is required for extensive branching and therefore for formation of these fine roots (Rasmussen et al., 2013; Jiu et al., 2022). Our data suggest there exists a molecular strategy for frost hardiness that prefers a desensitized strigoactone pathway that is less sensitive to internal signaling molecules such as the GOLVEN peptides. A more robust ability to reform fine roots and indeed Nitrogen fixing nodules would provide a survival advantage in regions that experience a higher frequency of frost days.

In summary, through a combination of GWAS and expression studies, we identified an interaction between the strigolactone and GOLVEN signaling pathways. This approach revealed distinct targets not identified by expression studies alone, underscoring the efficacy of GWAS in detecting downstream components of peptide signaling. Future investigation of the remaining 73 SNPs associated with legume nodulation and root architecture traits uncovered in this study will shed light on the molecular components and regulatory networks shaping beneficial legume-microbe interactions and root development.

## METHODS

### Plant material, nodulation assays and growth conditions

*Medicago truncatula* ecotype Jemalong A17 or R108 were used in this study. All *Tnt1* mutants are in the R108 background. *Tnt1* lines isolated include *d14-1* (NF18262). List of HapMap line accessions is provided in Supplementary Table 2. Seeds were scarified with concentrated H_2_SO_4_, and surface sterilized with undiluted household bleach (Clorox) at 8% sodium hypochlorite and sown on 1% water agarose (Life Technologies Catalog: 16500100) plates (Boschiero et al., 2019).

Seeds were stratified at 4°C for three days in dark prior to overnight germination at 24oC and then transferred onto agarose plates containing B&D nutrients plus 0.5 mM KNO_3_ with and without peptides. Eight to ten seedlings per line were placed on each plate between sterile filter paper sheets for all experiments except the GWAS screen in which we placed three seedlings per plate. Three days post transfer, seedlings from either treatment were inoculated with *S. meliloti* Rm 2011 or a rhizobia mixture of *S*. *meliloti*-KH46c and *S*.*medicae*-WSM419 to a final OD_600_ of 0.05. Phenotypic traits were scored under a microscope 4x lens objective (Leica S7). For experiments in soil, overnight germinated seedlings were transferred to a 2:1 mixture of turface:vermiculite. Plants were watered B&D solution containing six millimolar nitrogen before inoculation with rhizobia at seven days post germination. Post Inoculation, plants were subsequently watering with B&D nutrient media supplemented with 0.5 mM Potassium Nitrate. All experiments were conducted in a controlled environment chamber at 24°C under 16 hours light, eight hours dark conditions.

### Cloning of promoters and CDS for hairy root

The promoter sequences of *MtDWARF14* were integrated upstream of the β-Galactouronidase (GUS) gene in the MU06 vector, which contains a DsRed selection marker, using the Gibson assembly method. The *MtGLV10* and *MtD14* genes were cloned into the MU41 vector using Gibson assembly. Prior to introducing them into *M. truncatula* hairy roots using *Agrobacterium rhizogenes* Arqua1, all clones were verified using Sanger sequencing.

### Hairy root transformation

A strain of *A. rhizogenes* Arqua1 resistant to streptomycin was introduced with desired constructs that had an AtUBI:dsred selection tag. They were then grown on Luria-Bertani medium agar plates with appropriate antibiotics at 28°C for two days. Using clean spreaders, agrobacteria were collected from the plates and dissolved in 500-700 µL of sterile water. The root meristem of 1-day old germinated seedlings were removed by cutting off the root tips, and the cut section was then immersed in the previously mentioned bacterial mixture. These seedlings were then placed on modified Fahraeus medium plates and cultivated for two weeks at 24°C under 16 hours of daylight and 8 hours of nighttime conditions. Using an Olympus SZX microscope, transgenic calluses showing dsRed expression were identified and subsequently transferred to soil.

### Histochemical localization of GUS and β-Gal staining

For X-gluc staining, X-GlcA ((5-bromo-4-chloro-3-indolyl-beta-D-glucuronic acid, Goldbio) in DMF (Dimethyl formamide)) was added to 50 ml of phosphate buffer (100 mM phosphate buffer saline with 100 mM Na_2_HPO_4_, NaH_2_PO_4_ each) and 50 mM K_4_FeCN_6_ and K_3_FeCN_6_ each, 0.5M EDTA and 10% Triton X-100. X-gluc was added to a final concentration of 100 mg/100 mL buffer. Harvested root tissue were vacuum infiltrated with the X-Gluc solution for 10 minutes and then incubated at 37 °C in dark for varying time periods. Roots of *MtD14* were stained for 6 hours for each time point. At 28 dpi, nodules were excised and cleared with 1/20 strength bleach overnight and imaged.

Samples were triple-rinsed with phosphate buffer before using X-gal (Goldbio) for rhizobia staining. Before applying the X-gal (5-bromo-4-chloro-3-indolyl-β-D-galactopyranoside) stain, hairy roots were pre-treated with 2.5% glutaraldehyde using vacuum infiltration for a duration of ten minutes and then left to incubate for an additional hour. Afterwards, the samples were washed using Z-buffer (comprising 100 mM each of Na2HPO4 and NaH2PO4, 10 mM of Potassium chloride, and 1 mM of Magnesium chloride). They were then immediately placed in a Z-buffer-based X-gal solution (with 50 mM each of K4FeCN6 and K3FeCN6, along with 4% X-gal in DMF). After another round of vacuum infiltration, the samples were incubated in the dark to incubate overnight and were photographed the next day.

### Genome Wide Association Studies

Genome wide association studies (GWAS) were performed on 171 *M. truncatula* genotypes from the HapMap panel using Mt5.0 SNP markers (https://medicago.toulouse.inra.fr/MtrunA17r5.0-ANR/) implemented with heterozygous SNPs from the Mt4.0 SNP marker file (http://www.medicagohapmap.org/), which were missing in the original Mt.5.0 SNP file. SNPs were filtered with a minimum count of 145 (85% of the population) and minor allele frequency of 3%. After filtering, 1,795,542 SNPs remained and were used in the GWAS. Kinship matrix was calculated in TASSEL 5.2.30 using the centered IBS approach. Population structure (Q value) analysis was performed in Structure 2.3.4 using randomly selected 10,000 SNPs across eight chromosomes. The optimal GWAS model for each trait was selected based on Bayesian information criterion (BIC) evaluation using R packages “coxme” and “MuMIn” following McKown et al (2014) (McKown et al., 2014). For each trait, the model generating the lowest BIC value was adopted in the GWAS analysis. Models evaluated in the BIC calculations included General Linear Model (GLM), GLM with Q correction, Mixed Linear Model (MLM), and MLM with Q correction. After performing GWAS analysis in TASSEL, false discovery rates (FDR) of SNP associative p values were calculated using the “fdr” method in R. The cutoff threshold for significant SNP selection was FDR at 0.05.

### Hormone Treatments and Nodule excision

Peptides synthesis was carried out by Pepscan and Austin Chemicals, Inc. For all peptide and hormone treatments, overnight germinated *M. truncatula* seedlings were transferred onto 1% Agarose in B & D media and grown for three days at 24°C. Sixty to seventy seedlings were transferred to sterile water (pH adjusted to 6.8) containing the desired peptide/hormone and treatment allowed to proceed for three hours. Post treatment, roots were excised from 20 seedlings per replicate per biological replicate and shoots discarded. Solution designated as Full-N constituted six mM Nitrogen (Final concentration 0.5 mM KH_2_PO_4_, 0.25 mM K_2_SO_4_, 0.25 mM MgSO_4_, .01 mM Fe-citrate, 1 mM CaCl_2_, 2 mM KNO_3_, 2 mM NH_4_NO_3_, pH 6.8) while Low-N solution contained a limited amount of Nitrogen, prepared without any NH_4_NO_3_ and only 0.5 mM KNO_3_.

### RNA Extraction, complimentary DNA synthesis and quantitative PCR

Total RNA was extracted using Trizol Reagent (Life Technologies) following the manufacturer’s recommendations (Invitrogen GmbH, Karlsruhe, Germany), digested with RNase free DNase1 (Ambion Inc., Houston, TX) and column purified with RNeasy MinElute CleanUp Kit (Qiagen). RNA was quantified using a Nanodrop Spectrophotometer ND-100 (NanoDrop Technologies, Wilington, DE). RNA integrity was assessed on an Agilent 2100 BioAnalyser and RNA 6000 Nano Chips (Agilient Technologies, Waldbronn, Germany). First-strand complementary DNA was synthesized by priming with oligo-dT_20_ (Qiagen, Hilden, Germany), using Super Script Reverse Transcriptase III (Invitrogen GmbH, Karlsruhe, Germany) following manufacturer’s recommendations. Primer Express V3.0 software was used for primer design. qPCR reactions were carried out in an QuantStudio7 (ThermoFisher Scientific Inc.). Five microliters reactions were performed in an optical 384-well plate containing 2.5 μL SYBR Green Power Master Mix reagent (Applied Biosystems), 15 ng cDNA and 200 nM of each gene-specific primer. Transcript levels were normalized using the geometric mean of two housekeeping genes, *MtUBI* (Medtr3g091400) and *MtPTB* (Medtr3g090960). Three biological replicates were included and displayed as relative expression values. Primer sequences are provided in **Supplementary Table 4**.

### Imaging and Root scans

Traits were scored under a 4x objective lens in a Leica S7 microscope. Whole mount nodules were imaged after a 5% overnight bleach treatment using an Olympus SZX12 stereomicroscope (Olympus) equipped with a Nikon DXM1200C digital camera (Nikon Instruments). GUS staining images were taken with Olympus BX41 microscope equipped with an Olympus DP72 camera. Roots were scanned using an Epson 12000 XL scanner with a large transparency unit using acrylic scanning trays doi.org/10.5281/zenodo.4122422) and root length traits extracted using RhizoVision (Seethepalli et al., 2020; Seethepalli et al., 2021).

### Statistical Analysis

All statistical analyses were performed using GraphPad Prism 8 and tests selected therein. A two-sided Student’s t-test was used for comparison between genotypes or treatments. For multiple genotypes one-way analysis of variance tests were performed followed by post-hoc statistical tests as mentioned.

### Figure Preparation and R packages used

Figures were prepared using Adobe Illustrator Creative cloud. Images were edited using Adobe Photoshop and FIJI. Graphs were prepared using GraphPad Prism 8. Phylogenetic tree was prepared using Mega X and the FigTree Application. R packages used in this study; Rain plots; PCA Analysis (FactoMineR”), Bubble plot (https://rdrr.io/cran/GOplot/man/GOBubble.html). A correlation matrix between the root traits was computed as the pairwise Pearson correlation coefficients using the cor function in RStudio version 4.1.3. The R Corrplot package was utilized to visualize the distribution of correlation as a correlogram. The correlation coefficients were plotted for both fold change upon treatment as well as control samples, respectively. All default plots were edited using Adobe Illustrator for clarity.

## Supporting information

GWAS traits and associated statistically significant SNPs found in this study.

---

**Supplementary Table 1:** GWAS traits and associated statistically significant SNPs found in this study.

**Supplementary Table 2:** Genes within 100Kb of SNPs identified in this study.

**Supplementary Table 3:** Expression of Strigolactone related genes upon GLV10 application.

**Supplementary Table 4:** Raw values for each trait and HapMap line used in this study.

**Supplementary Table 5:** Pearson Correlation amongst traits in 131 Medicago HapMap accessions.

| Trait Number | Trait Name | Marker | Position | Allele | Location | pValue | qValue |  |
| --- | --- | --- | --- | --- | --- | --- | --- | --- |
| Trait 12 | NodNum | Chr1_13463336 | 13531246 |  | Intergenic | 3.4E-07 | 0.041 |  |
| Trait 12 | NodNum | Chr1_25459986 | 29437003 | C/A | Intron | 6.8E-09 | 0.006 |  |
| Trait 12 | NodNum | Chr1_25460966 | 29437983 | C/T | Intron | 1.5E-07 | 0.035 |  |
| Trait 12 | NodNum | Chr1_25462303 | 29439320 | C/T | Intron | 3.9E-07 | 0.044 |  |
| Trait 12 | NodNum | Chr2_31981499 | 37998699 |  | Intergenic | 4.2E-09 | 0.006 | Shared With Nodule Density |
| Trait 12 | NodNum | Chr2_36636572 | 42733785 | A/G | Exon | 2.1E-07 | 0.035 | Shared With Nodule Density |
| Trait 12 | NodNum | Chr3_23083763 | 26305255 | G/A | Intergenic | 5.6E-07 | 0.050 | Shared With Nodule Density |
| Trait 12 | NodNum | Chr3_35888202 | 39352531 |  | repeat_region | 1.9E-07 | 0.035 |  |
| Trait 12 | NodNum | Chr3_53916623 | 57339100 | G/C | repeat_region | 4.6E-07 | 0.049 |  |
| Trait 12 | NodNum | Chr4_42751410 | 50701554 | C/G | Exon | 2.1E-07 | 0.035 | Shared With Nodule Density |
| Trait 12 | NodNum | Chr4_44470744 | 52458276 | A/C | repeat_region | 2.6E-07 | 0.036 |  |
| Trait 12 | NodNum | Chr4_46642725 | 54753554 | G/A | repeat_region | 1.3E-07 | 0.035 |  |
| Trait 12 | NodNum | Chr5_20095271 | 20021305 | C/T | repeat_region | 1.9E-07 | 0.035 |  |
| Trait 12 | NodNum | Chr6_24591040 | 32546885 | A/T | Intergenic | 5.1E-07 | 0.049 |  |
| Trait 12 | NodNum | Chr6_35125786 | 42729136 | C/A | Intron | 1.4E-07 | 0.035 | Shared With Nodule Density |
| Trait 12 | NodNum | Chr6_35167950 | 42763851 | C/T | Intron | 1.7E-07 | 0.035 | Shared With Nodule Density |
| Trait 12 | NodNum | Chr8_11825410 | 11869212 | C/A | Intergenic | 1.4E-07 | 0.035 |  |
| Trait 12 | NodNum | Chr8_11825415 | 11869217 | G/C | Intergenic | 3.2E-07 | 0.041 | Shared With Nodule Density |
| Trait 12 | NodNum | Chr8_11861896 | 11936701 | C/T | Intron | 2.5E-07 | 0.036 | Shared With Nodule Density |
| Trait 12 | NodNum | Chr8_22195415 | 24465364 | T/C | Intron | 5.2E-07 | 0.049 |  |
| Trait 15 | NodDensity | Chr1_5202353 | 5255399 | T/C | 5' UTR | 5.6E-07 | 0.046 |  |
| Trait 15 | NodDensity | Chr3_2999343 | 3012769 | C/A | Exon | 1.4E-07 | 0.021 |  |
| Trait 15 | NodDensity | Chr4_46764973 | 54876129 | G/T | repeat_region | 3.5E-08 | 0.021 |  |
| Trait 15 | NodDensity | Chr4_51063347 | 59185345 | A/G | Intron | 1.2E-07 | 0.021 |  |
| Trait 15 | NodDensity | Chr4_51063378 | 59185376 | C/T | Intron | 1.1E-07 | 0.021 |  |
| Trait 15 | NodDensity | Chr4_51091479 | 59213477 | G/A | Exon | 8.8E-08 | 0.021 |  |
| Trait 15 | NodDensity | Chr7_24950580 | 31760798 | C/T | Intergenic | 1.3E-07 | 0.021 |  |
| Trait 15 | NodDensity | Chr8_21491467 | 21432873 | C/A | repeat_region | 2.5E-07 | 0.026 |  |
| Trait 15 | NodDensity | Chr8_22065532 | 22009539 | C/T | repeat_region | 9.8E-09 | 0.017 |  |
| Trait 15 | NodDensity | Chr8_32266780 | 34735229 | G/A | 3' UTR | 1.4E-07 | 0.021 |  |
| Trait 15 | NodDensity | scaffold0034_49163 | 22802642 | C/T | - | 1.6E-07 | 0.021 |  |
| Trait 15 | NodDensity | scaffold0190_11392 | 22435849 | G/C | - | 2.2E-07 | 0.026 |  |
| Trait 15 | NodDensity | scaffold0324_11411 | 24426226 | T/G | - | 5.4E-07 | 0.046 |  |
| Trait 15 | NodDensity | scaffold0598_5040 | 24382238 | A/G | - | 4.1E-07 | 0.037 |  |
| Trait 16 | RelNodPos | Chr2_22840616 | 22070192 | A/T | repeat_region | 9.5E-08 | 0.028 |  |
| Trait 16 | RelNodPos | Chr4_52006663 | 60132495 | C/G | repeat_region | 7.4E-09 | 0.013 |  |
| Trait 16 | RelNodPos | Chr4_6589625 | 7082288 | A/C | Exon | 4.9E-08 | 0.026 |  |
| Trait 16 | RelNodPos | Chr7_26120165 | 32861704 | C/T | repeat_region | 7.4E-08 | 0.026 |  |
| Trait 16 | RelNodPos | Chr8_17266814 | 21075727 | C/T | repeat_region | 2.1E-07 | 0.047 |  |
| Trait 16 | RelNodPos | Chr8_22282343 | 24545296 | A/C | repeat_region | 2.6E-07 | 0.047 |  |
| Trait 16 | RelNodPos | Chr8_22283375 | 24546328 | C/G | repeat_region | 2.4E-07 | 0.047 |  |
| Trait 16 | RelNodPos | Chr8_22289779 | 24552732 | G/A | repeat_region | 7.4E-08 | 0.026 |  |
| Trait 16 | RelNodPos | Chr8_22311333 | 24573418 | C/T | repeat_region | 1.2E-07 | 0.029 |  |
| Trait 16 | RelNodPos | Chr8_26113721 | 28615214 | C/T | repeat_region | 4.3E-08 | 0.026 |  |
| Trait 18 | NodZone | Chr3_14180965 | 14083424 | G/A | repeat_region | 6.3E-08 | 0.046 |  |
| Trait 18 | NodZone | Chr3_17458306 | 19185759 | T/C | repeat_region | 1.8E-09 | 0.003 |  |
| Trait 18 | NodZone | Chr8_3214636 | 3089447 | C/T | Intergenic | 7.6E-08 | 0.046 |  |
| Trait 2 | LRNprim | Chr3_53580691 | 57001431 | T/C | repeat_region | 1.8E-09 | 0.003 |  |
| Trait 4 | LRNelong | Chr1_20967038 | 24548473 |  | repeat_region | 1.3E-07 | 0.020 |  |
| Trait 4 | LRNelong | Chr2_17421352 | 17456761 |  | repeat_region | 6.7E-08 | 0.013 |  |
| Trait 4 | LRNelong | Chr2_18430856 | 18528899 | C/A | repeat_region | 2.8E-07 | 0.031 |  |
| Trait 4 | LRNelong | Chr2_18430864 | 18528907 | T/G | repeat_region | 2.9E-07 | 0.031 |  |
| Trait 4 | LRNelong | Chr2_38585840 | 44702198 | T/A | repeat_region | 7.0E-09 | 0.004 |  |
| Trait 4 | LRNelong | Chr3_42094632 | 45554638 | T/C | Gene - Exon | 3.5E-07 | 0.033 |  |
| Trait 4 | LRNelong | Chr3_7792838 | 8285386 | C/T | repeat_region | 4.7E-09 | 0.004 |  |
| Trait 4 | LRNelong | Chr4_14724836 | 16776663 |  | repeat_region | 2.0E-07 | 0.025 |  |
| Trait 4 | LRNelong | Chr4_22331217 | 30540815 | C/T | Exon | 8.2E-08 | 0.015 |  |
| Trait 4 | LRNelong | Chr4_51684839 | 59807893 | C/A | repeat_region | 8.6E-09 | 0.004 |  |
| Trait 4 | LRNelong | Chr4_55376508 | 63553388 | T/C | repeat_region | 5.5E-08 | 0.012 |  |
| Trait 4 | LRNelong | Chr5_15177298 | 15088738 | G/A | Exon | 2.0E-08 | 0.007 |  |
| Trait 4 | LRNelong | Chr5_24008362 | 25808752 | A/C | repeat_region | 3.9E-08 | 0.010 |  |
| Trait 4 | LRNelong | Chr5_6050116 | 6057978 | A/C | Intergenic | 3.0E-08 | 0.009 |  |
| Trait 4 | LRNelong | Chr6_15007206 | 16078787 | A/T | repeat_region | 5.6E-07 | 0.050 |  |
| Trait 4 | LRNelong | Chr7_14322836 | 17797531 |  | repeat_region | 2.3E-07 | 0.028 |  |
| Trait 4 | LRNelong | Chr7_18719061 | 25168558 | G/A | 3' UTR | 1.6E-07 | 0.022 |  |
| Trait 4 | LRNelong | Chr7_18719062 | 25168559 | C/T | 3' UTR | 1.3E-07 | 0.020 |  |
| Trait 4 | LRNelong | Chr8_14675398 | 15046467 | G/A | repeat_region | 8.0E-09 | 0.004 |  |
| Trait 4 | LRNelong | Chr8_43680693 | 47813927 | C/T | repeat_region | 3.4E-07 | 0.033 |  |
| Trait 9 | totalLRdensity | Chr6_17989136 | 19480800 | C/A | Exon | 2.0E-08 | 0.009 |  |
| Trait 9 | totalLRdensity | Chr6_17989170 | 19480834 | G/A | Exon | 7.3E-09 | 0.006 |  |
| Trait 9 | totalLRdensity | Chr6_17989303 | 19480967 | C/G | Intergenic | 1.1E-08 | 0.006 |  |
| Trait 9 | totalLRdensity | Chr6_17989326 | 19480990 | T/G | Intergenic | 7.1E-09 | 0.006 |  |
| Trait 9 | totalLRdensity | Chr6_17990127 | 19481791 | G/A | Intergenic | 4.5E-08 | 0.016 |  |
| Trait 9 | totalLRdensity | Chr7_29068050 | 35770510 | G/A | Intergenic | 1.6E-07 | 0.048 |  |

## SUPPLEMENTARY FIGURES

**Figure S1:**
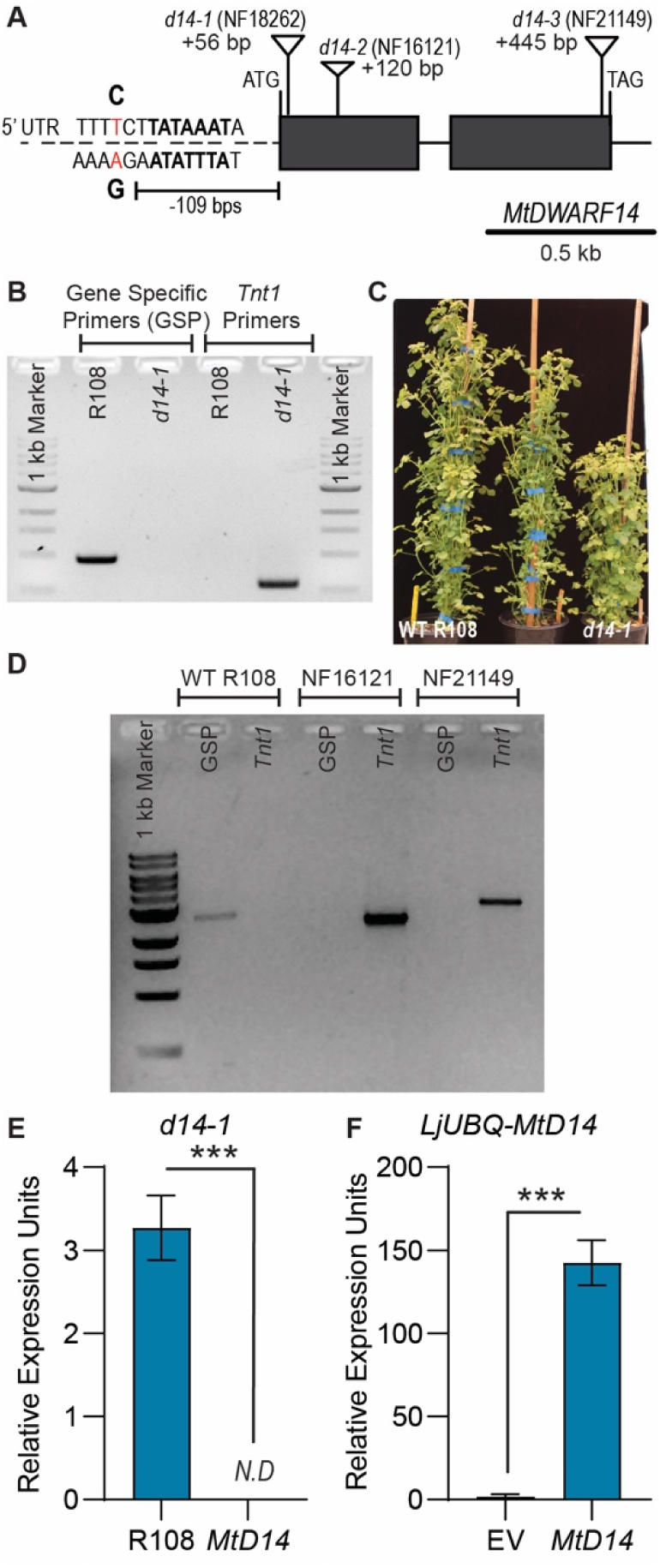
Characterization of lines used in this study. Gene structure of the MtDWARF15 gene indicating positions of Tnt1 inserts and location of the SNP identified using GWAS. Gel image showing presence of the Tnt1 insert in d14-1 (NF18262) lines compared to WT R108. Three-month-old Medicago WT R108 plants (left) compared to a dwarf d14-1 mutant (right). Gel image showing presence of the Tnt1 insert in d14-2 (NF16121) and d14-3 (NF21149) lines compared to WT R108. Quantitative PCR comparing transcript levels of MtDWARF14 in the WT R108 background and d14-1 lines. Three biological replicated each with 20 seedling roots were used for RNA extraction. Student’s t-test ***p<0.001 Quantitative PCR comparing transcript levels of MtDWARF14 in transgenic empty vector lines compared to MtDWARF14 over-expressing lines. At least 3-5 lines per biological replicate were used. Student’s t-test ***p<0.001. Data normalized to two housekeeping genes MtUBQ and MtPTB.

**Figure S2:**
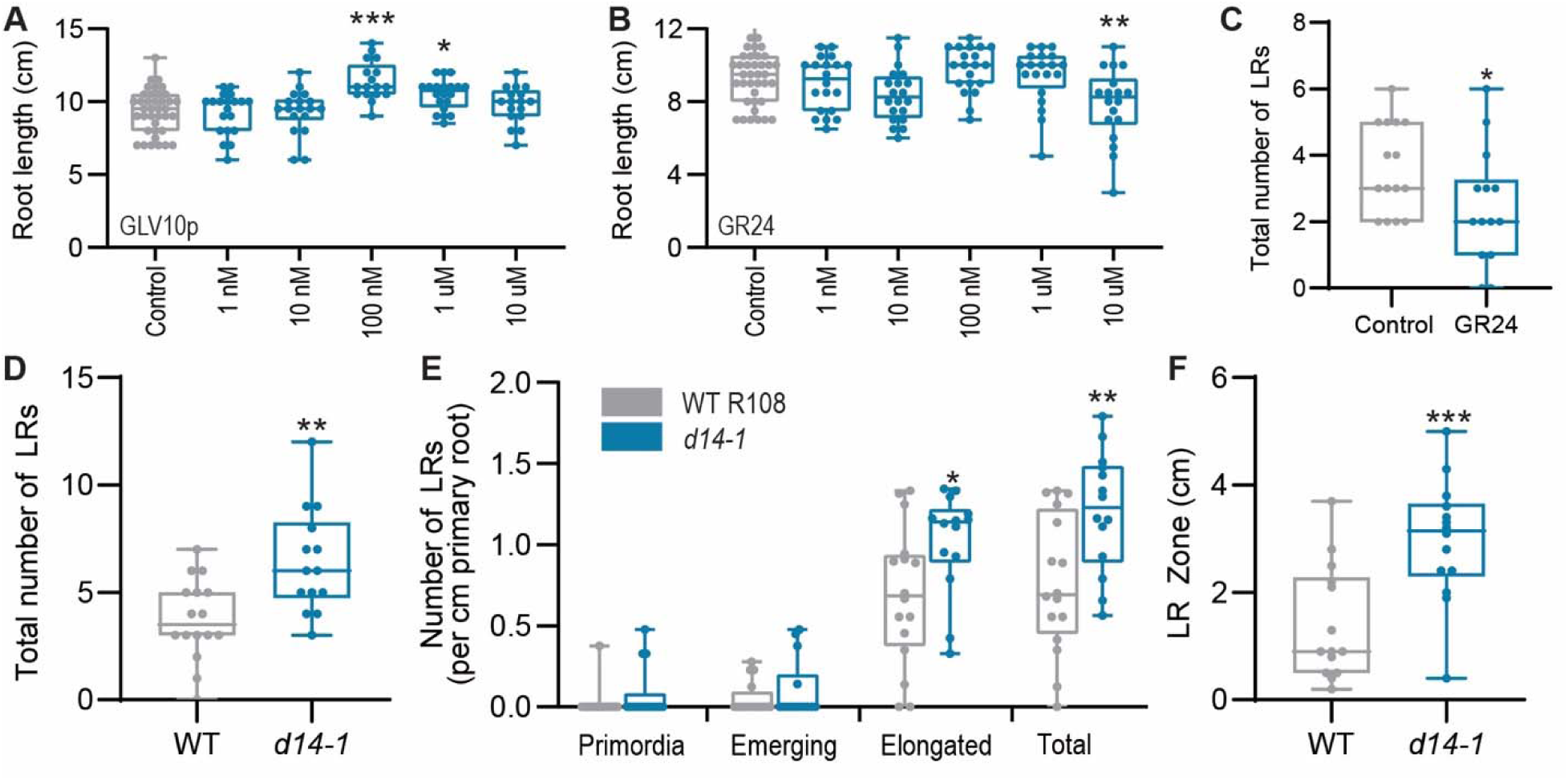
Lateral root traits upon GR24 application and in the d14-1 null mutant. Primary root length change in *M. truncatula* Jemalong A17 upon application of GLV10 peptide A. and B. synthetic strigolactone analogue GR24 at concentrations indicated. For each treatment n=13-20. Asterisks indicate ANOVA protected Dunnett’s comparison test where *p<0.05, **p<0.01, ***p<0.001. C. Boxplot showing total number of lateral roots upon application or GR24 compared to the solvent control. D. Number of lateral roots in d14-1 mutant lines compared to control WT R108 seedling roots seven days post germination. Student’s t-test **p<0.01. E. Number of lateral roots per cm of primary root in d14-1 mutant lines compared to control R108 seedling roots seven days post germination. Student’s t-test *p<0.05, **p<0.01. F. Length of lateral root zone in d14-1 mutant lines compared to control R108 seedling roots seven days post germination. Student’s t-test *p<0.05.

**Supplementary Figure S3:**
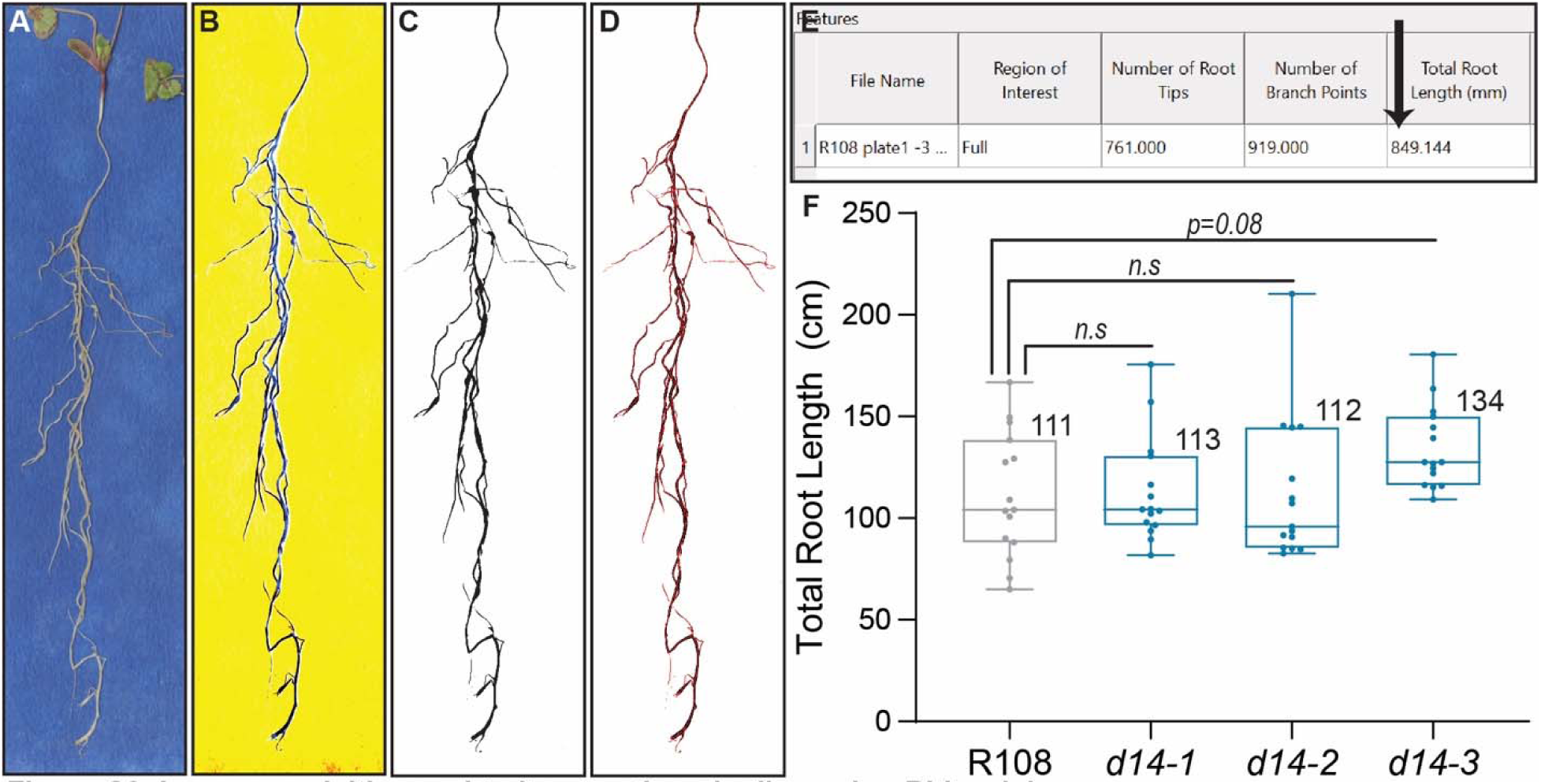
Image acquisition and trait extraction pipeline using Rhizovision. A. Raw image of *M. truncatula* roots scanned against a blue background. B. Edited image in photoshop after inverting colors and altering color ‘Curves’ to obtain a solid color background. C. Image processed by Rhizovision using default settings. D. Output image showing area of the root selected by the software to measure root length. E. Screenshot showing traits considered in the output provided by Rhizovision. F. Box plot showing total root length of d14 mutant alleles compared to wild type R108. Sample size is 15 for all groups. Average values are shown on the shoulder of each box plot. Values were tested using ordinary one way ANOVA.

**Figure S4.**
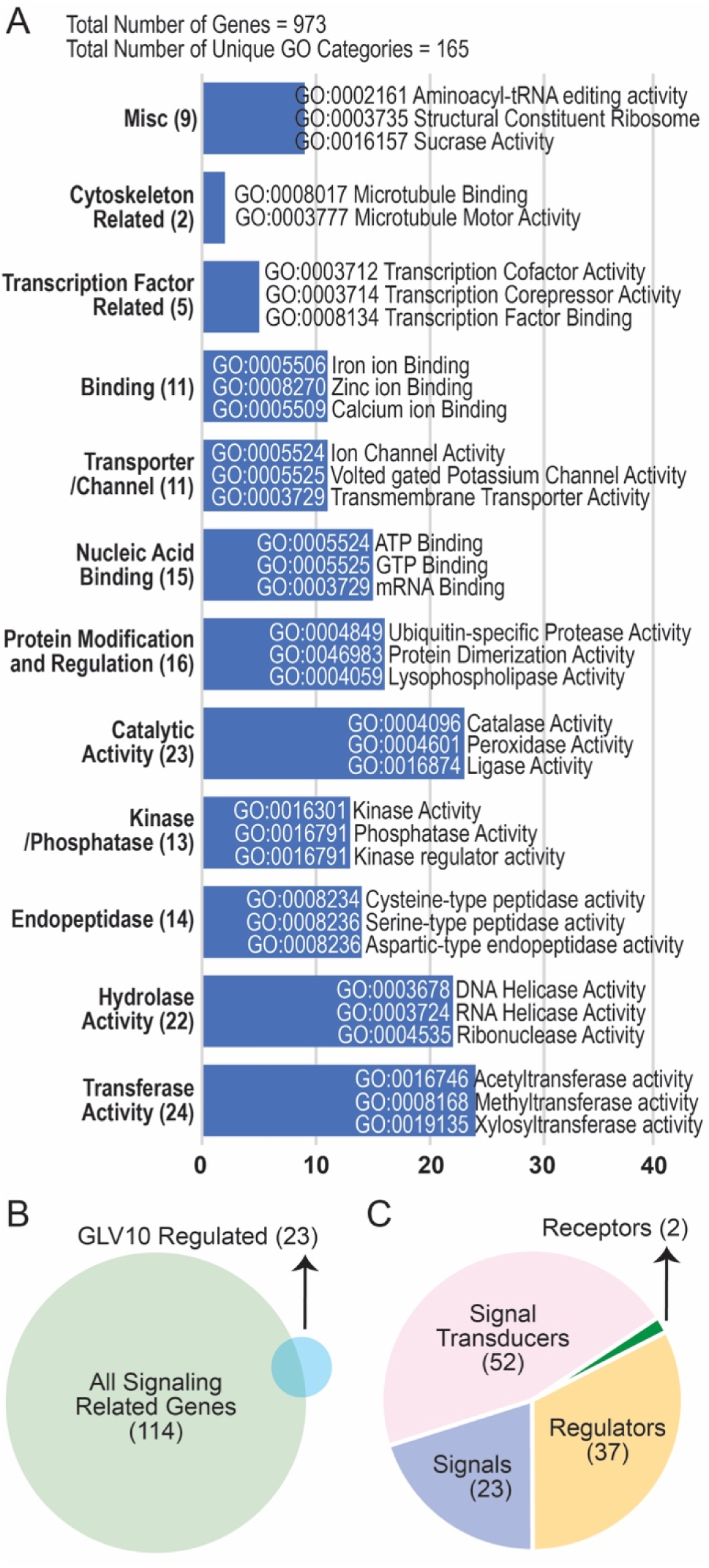
Gene Ontology Classification of endidate genes with 100 kb of SNPs identified in this study. A. Histogram showing clusters of GO categories most highly enriched. Numbers in brackets indicate the number of GO terms. B. Venn Diagram showing total number of signaling related genes and percent overlap with those transcriptionally regulated by the peptide GOLVEN10. C. Pie chart showing sub categories of signaling genes identified in this study.

## ACKNOWLEDGEMENTS

Authors gratefully acknowledge the help of Lynne Jacobs in the greenhouse. We would like to thanks Dr(s). Jimena Guerrero and Stephanie Manel for kindly sharing their raw data on climatic variables associated with regions where the HapMap lines were collected. This research was funded by the award National Science Foundation Award #1444549 to Wolf Schieble and Michael Udvardi and by the United States Department of Agriculture grants 2022-38821-37353 and National Science Foundation Award #2217830 to Sonali Roy.

## AUTHOR CONTRIBUTIONS

S.R, S.Z, I.T.J, D.J, B.S, X. C performed experiments, S.R, Y.K, J. W analyzed data, S.R, W.R.S, M.K.U conceptualized this study, M.K.U, W.R.S, J.D.M, supervised the research, S.R, M.K.U wrote the manuscript.

## REFERENCES

1. Akiyama, K., Matsuzaki, K., and Hayashi, H. (2005). Plant sesquiterpenes induce hyphal branching in arbuscular mycorrhizal fungi. Nature 435, 824–827.

2. Ambroise, V., Legay, S., Guerriero, G., Hausman, J.-F., Cuypers, A., and Sergeant, K. (2020). The roots of plant frost hardiness and tolerance. Plant and Cell Physiology 61, 3–20.

3. Banasiak, J., Borghi, L., Stec, N., Martinoia, E., and Jasiński, M. (2020). The Full-Size ABCG Transporter of Medicago truncatula Is Involved in Strigolactone Secretion, Affecting Arbuscular Mycorrhiza. Front Plant Sci 11, 18.

4. Bensmihen, S. (2015). Hormonal control of lateral root and nodule development in legumes. Plants 4, 523–547.

5. Bonhomme, M., Bensmihen, S., André, O., Amblard, E., Garcia, M., Maillet, F., Puech-Pagès, V., Gough, C., Fort, S., and Cottaz, S. (2021). Distinct genetic basis for root responses to lipo-chitooligosaccharide signal molecules from distinct microbial origins. Journal of Experimental Botany.

6. Boschiero, C., Lundquist, P.K., Roy, S., Dai, X., Zhao, P.X., and Scheible, W.R. (2019). Identification and Functional Investigation of Genome-Encoded, Small, Secreted Peptides in Plants. Curr Protoc Plant Biol 4, e20098.

7. Breakspear, A., Liu, C., Roy, S., Stacey, N., Rogers, C., Trick, M., Morieri, G., Mysore, K.S., Wen, J., Oldroyd, G.E.D., Downie, J.A., and Murray, J.D. (2014). The root hair “Infectome” of *Medicago truncatula* uncovers changes in cell cycle genes and reveals a requirement for auxin signaling in rhizobial infection. The Plant Cell 26, 4680–4701.

8. Brooks, E.G., Elorriaga, E., Liu, Y., Duduit, J.R., Yuan, G., Tsai, C.-J., Tuskan, G.A., Ranney, T.G., Yang, X., and Liu, W. (2023). Plant promoters and terminators for high-precision bioengineering. BioDesign Research 5, 0013.

9. Bürger, M., and Chory, J. (2020). The many models of strigolactone signaling. Trends in plant science 25, 395–405.

10. Chen, Z., Lancon-Verdier, V., Le Signor, C., She, Y.-M., Kang, Y., and Verdier, J. (2021a). Genome-wide association study identified candidate genes for seed size and seed composition improvement in M. truncatula. Scientific Reports 11, 4224.

11. Chen, Z., Ly Vu, J., Ly Vu, B., Buitink, J., Leprince, O., and Verdier, J. (2021b). Genome-wide association studies of seed performance traits in response to heat stress in Medicago truncatula uncover miel1 as a regulator of seed germination plasticity. Frontiers in plant science 12, 673072.

12. Chevalier, F., Nieminen, K., Sánchez-Ferrero, J.C., Rodríguez, M.L., Chagoyen, M., Hardtke, C.S., and Cubas, P. (2014). Strigolactone promotes degradation of DWARF14, an α/β hydrolase essential for strigolactone signaling in Arabidopsis. The Plant Cell 26, 1134–1150.

13. Demirjian, C., Vailleau, F., Berthomé, R., and Roux, F. (2023). Genome-wide association studies in plant pathosystems: success or failure? Trends in plant science 28, 471–485.

14. Epstein, B., Burghardt, L.T., Heath, K.D., Grillo, M.A., Kostanecki, A., Hämälä, T., Young, N.D., and Tiffin, P. (2023). Combining GWAS and population genomic analyses to characterize coevolution in a legume_-_rhizobia symbiosis. Molecular ecology 32, 3798–3811.

15. Fernandez, A., Drozdzecki, A., Hoogewijs, K., Vassileva, V., Madder, A., Beeckman, T., and Hilson, P. (2015). The GLV6/RGF8/CLEL2 peptide regulates early pericycle divisions during lateral root initiation. Journal of experimental botany 66, 5245–5256.

16. Fernandez, A.I., Vangheluwe, N., Xu, K., Jourquin, J., Claus, L.A.N., Morales-Herrera, S., Parizot, B., De Gernier, H., Yu, Q., Drozdzecki, A., Maruta, T., Hoogewijs, K., Vannecke, W., Peterson, B., Opdenacker, D., Madder, A., Nimchuk, Z.L., Russinova, E., and Beeckman, T. (2020). GOLVEN peptide signalling through RGI receptors and MPK6 restricts asymmetric cell division during lateral root initiation. Nature Plants 6, 533–543.

17. Foo, E., and Davies, N.W. (2011). Strigolactones promote nodulation in pea. Planta 234, 1073–1081.

18. Franssen, H.J., Xiao, T.T., Kulikova, O., Wan, X., Bisseling, T., Scheres, B., and Heidstra, R. (2015). Root developmental programs shape the Medicago truncatula nodule meristem. Development 142, 2941–2950.

19. Grace, M.L., Chandrasekharan, M.B., Hall, T.C., and Crowe, A.J. (2004). Sequence and spacing of TATA box elements are critical for accurate initiation from the beta-phaseolin promoter. J Biol Chem 279, 8102–8110.

20. Guerrero, J., Andrello, M., Burgarella, C., and Manel, S. (2018). Soil environment is a key driver of adaptation in Medicago truncatula: new insights from landscape genomics. New Phytologist 219, 378–390.

21. Herrbach, V., Remblière, C., Gough, C., and Bensmihen, S. (2014). Lateral root formation and patterning in *Medicago truncatula*. Journal of Plant Physiology 171, 301–310.

22. Hirsch, A.M., Larue, T.A., and Doyle, J. (1997). Is the legume nodule a modified root or stem or an organ sui generis? Critical Reviews in Plant Sciences 16, 361–392.

23. Hofhuis, H., Laskowski, M., Du, Y., Prasad, K., Grigg, S., Pinon, V., and Scheres, B. (2013). Phyllotaxis and rhizotaxis in Arabidopsis are modified by three PLETHORA transcription factors. Current Biology 23, 956–962.

24. Ji, H., Xiao, R., Lyu, X., Chen, J., Zhang, X., Wang, Z., Deng, Z., Wang, Y., Wang, H., and Li, R. (2022). Differential light-dependent regulation of soybean nodulation by papilionoid-specific HY5 homologs. Current Biology 32, 783–795. e785.

25. Jiang, L., Matthys, C., Marquez-Garcia, B., De Cuyper, C., Smet, L., De Keyser, A., Boyer, F.-D., Beeckman, T., Depuydt, S., and Goormachtig, S. (2016). Strigolactones spatially influence lateral root development through the cytokinin signaling network. Journal of Experimental Botany 67, 379–389.

26. Jiu, S., Xu, Y., Wang, J., Haider, M.S., Xu, J., Wang, L., Wang, S., Li, J., Liu, X., and Sun, W. (2022). Molecular mechanisms underlying the action of strigolactones involved in grapevine root development by interacting with other phytohormone signaling. Scientia Horticulturae 293, 110709.

27. Johnson, D., and Rumbaugh, M. (1986). Field nodulation and acetylene reduction activity of high-altitude legumes in the western United States. Arctic and Alpine Research 18, 171–179.

28. Jourquin, J., Fernandez, A.I., Xu, K., Simura, J., Ljung, K., and Beeckman, T. (2022). GOLVEN peptides regulate lateral root spacing as part of a negative feedback loop on the establishment of auxin maxima. bioRxiv, 2022.2009. 2026.509595.

29. Jourquin, J., Fernandez, A.I., Wang, Q., Xu, K., Chen, J., Šimura, J., Ljung, K., Vanneste, S., and Beeckman, T. (2023). GOLVEN peptides regulate lateral root spacing as part of a negative feedback loop on the establishment of auxin maxima. Journal of Experimental Botany 74, 4031–4049.

30. Jullien, M., Ronfort, J., and Gay, L. (2021). How and When Does Outcrossing Occur in the Predominantly Selfing Species Medicago truncatula? Front Plant Sci 12, 619154.

31. Junior, M.d.A.L., Lima, A., Arruda, J., and Smith, D. (2005). Effect of root temperature on nodule development of bean, lentil and pea. Soil Biology and Biochemistry 37, 235–239.

32. Kang, Y., Torres_-_Jerez, I., An, Z., Greve, V., Huhman, D., Krom, N., Cui, Y., and Udvardi, M. (2019). Genome_-_ wide association analysis of salinity responsive traits in Medicago truncatula. Plant, Cell & Environment 42, 1513–1531.

33. Kiran, K., Ansari, S.A., Srivastava, R., Lodhi, N., Chaturvedi, C.P., Sawant, S.V., and Tuli, R. (2006). The TATA-box sequence in the basal promoter contributes to determining light-dependent gene expression in plants. Plant Physiol 142, 364–376.

34. Koltai, H. (2015). Cellular events of strigolactone signalling and their crosstalk with auxin in roots. Journal of experimental botany 66, 4855–4861.

35. Lee, T., Orvosova, M., Batzenschlager, M., Batista, M.B., Bailey, P.C., Mohd-Radzman, N.A., Gurzadyan, A., Stuer, N., Mysore, K.S., and Wen, J. (2024). Light-sensitive short hypocotyl genes confer symbiotic nodule identity in the legume Medicago truncatula. Current Biology 34, 825–840. e827.

36. Li, Q., Li, M., Zhang, D., Yu, L., Yan, J., and Luo, L. (2020). The peptide-encoding *MtRGF3* gene negatively regulates nodulation of *Medicago truncatula*. Biochemical and Biophysical Research Communications 523, 66–71.

37. Małolepszy, A., Mun, T., Sandal, N., Gupta, V., Dubin, M., Urbański, D., Shah, N., Bachmann, A., Fukai, E., and Hirakawa, H. (2016). The LORE 1 insertion mutant resource. The Plant Journal 88, 306–317.

38. Mashiguchi, K., Seto, Y., and Yamaguchi, S. (2021). Strigolactone biosynthesis, transport and perception. The Plant Journal 105, 335–350.

39. Matsuzaki, Y., Ogawa-Ohnishi, M., Mori, A., and Matsubayashi, Y. (2010). Secreted peptide signals required for maintenance of root stem cell niche in *Arabidopsis*. Science 329, 1065–1067.

40. McKown, A.D., Guy, R.D., Klápště, J., Geraldes, A., Friedmann, M., Cronk, Q.C., El_-_Kassaby, Y.A., Mansfield, S.D., and Douglas, C.J. (2014). Geographical and environmental gradients shape phenotypic trait variation and genetic structure in *Populus trichocarpa*. New Phytologist 201, 1263–1276.

41. Mohd-Radzman, N.A., Djordjevic, M.A., and Imin, N. (2013). Nitrogen modulation of legume root architecture signaling pathways involves phytohormones and small regulatory molecules. Frontiers in Plant Science 4, 385.

42. Muller, L.M., Flokova, K., Schnabel, E., Sun, X., Fei, Z., Frugoli, J., Bouwmeester, H.J., and Harrison, M.J. (2019). A CLE-SUNN module regulates strigolactone content and fungal colonization in arbuscular mycorrhiza. Nat Plants 5, 933–939.

43. Na, D., Son, H., and Gsponer, J. (2014). Categorizer: a tool to categorize genes into user-defined biological groups based on semantic similarity. BMC genomics 15, 1–11.

44. Nandety, R.S., Wen, J., and Mysore, K.S. (2023). Medicago truncatula resources to study legume biology and symbiotic nitrogen fixation. Fundamental Research 3, 219–224.

45. Pislariu, C.I., Murray, J.D., Wen, J., Cosson, V., Muni, R.R., Wang, M., Benedito, V.A., Andriankaja, A., Cheng, X., Jerez, I.T., Mondy, S., Zhang, S., Taylor, M.E., Tadege, M., Ratet, P., Mysore, K.S., Chen, R., and Udvardi, M.K. (2012). A Medicago truncatula tobacco retrotransposon insertion mutant collection with defects in nodule development and symbiotic nitrogen fixation. Plant Physiol 159, 1686–1699.

46. Rasmussen, A., Depuydt, S., Goormachtig, S., and Geelen, D. (2013). Strigolactones fine-tune the root system. Planta 238, 615–626.

47. Rellán-Álvarez, R., Lobet, G., and Dinneny, J.R. (2016). Environmental control of root system biology. Annual review of plant biology 67, 619–642.

48. Rey, T., Bonhomme, M., Chatterjee, A., Gavrin, A., Toulotte, J., Yang, W., André, O., Jacquet, C., and Schornack, S. (2017). The Medicago truncatula GRAS protein RAD1 supports arbuscular mycorrhiza symbiosis and Phytophthora palmivora susceptibility. Journal of Experimental Botany 68, 5871–5881.

49. Ristova, D., Giovannetti, M., Metesch, K., and Busch, W. (2018). Natural genetic variation shapes root system responses to phytohormones in Arabidopsis. Plant Journal 96, 468–481.

50. Roy, S., and Muller, L.M. (2022). A rulebook for peptide control of legume-microbe endosymbioses. Trends Plant Sci.

51. Roy, S., Liu, W., Nandety, R.S., Crook, A., Mysore, K.S., Pislariu, C.I., Frugoli, J., Dickstein, R., and Udvardi, M.K. (2020). Celebrating 20 years of genetic discoveries in legume nodulation and symbiotic nitrogen fixation. The Plant Cell 32, 15–41.

52. Roy, S., Torres-Jerez, I., Zhang, S., Liu, W., Schiessl, K., Jain, D., Boschiero, C., Lee, H.K., Krom, N., Zhao, P.X., Murray, J.D., Oldroyd, G.E.D., Scheible, W.R., and Udvardi, M. (2024). The peptide GOLVEN10 alters root development and noduletaxis in Medicago truncatula. Plant J.

53. Sauer, M., and Kleine-Vehn, J. (2019). PIN-FORMED and PIN-LIKES auxin transport facilitators. Development 146.

54. Schaefer, R.J., Michno, J.-M., Jeffers, J., Hoekenga, O., Dilkes, B., Baxter, I., and Myers, C.L. (2018). Integrating coexpression networks with GWAS to prioritize causal genes in maize. The Plant Cell 30, 2922–2942.

55. Schiessl, K., Lilley, J.L.S., Lee, T., Tamvakis, I., Kohlen, W., Bailey, P.C., Thomas, A., Luptak, J., Ramakrishnan, K., Carpenter, M.D., Mysore, K.S., Wen, J., Ahnert, S., Grieneisen, V.A., and Oldroyd, G.E.D. (2019). NODULE INCEPTION recruits the lateral root developmental program for symbiotic nodule organogenesis in *Medicago truncatula*. Current Biology 29, 3657–3668.e3655.

56. Seethepalli, A., Dhakal, K., Griffiths, M., Guo, H., Freschet, G.T., and York, L.M. (2021). RhizoVision Explorer: open-source software for root image analysis and measurement standardization. AoB Plants 13, plab056.

57. Seethepalli, A., Guo, H., Liu, X., Griffiths, M., Almtarfi, H., Li, Z., Liu, S., Zare, A., Fritschi, F.B., and Blancaflor, E.B. (2020). Rhizovision crown: An integrated hardware and software platform for root crown phenotyping. Plant Phenomics 2020.

58. Shi, W., and Zhou, W. (2006). Frequency distribution of TATA Box and extension sequences on human promoters. BMC bioinformatics 7, 1–12.

59. Stanton-Geddes, J., Paape, T., Epstein, B., Briskine, R., Yoder, J., Mudge, J., Bharti, A.K., Farmer, A.D., Zhou, P., and Denny, R. (2013). Candidate genes and genetic architecture of symbiotic and agronomic traits revealed by whole-genome, sequence-based association genetics in Medicago truncatula. PloS one 8, e65688.

60. Swarbreck, S.M., Mohammad-Sidik, A., and Davies, J.M. (2020). Common components of the strigolactone and karrikin signaling pathways suppress root branching in Arabidopsis. Plant Physiology 184, 18–22.

61. Tabata, R., Sumida, K., Yoshii, T., Ohyama, K., Shinohara, H., and Matsubayashi, Y. (2014). Perception of root-derived peptides by shoot LRR-RKs mediates systemic N-demand signaling. Science 346, 343–346.

62. Tadege, M., Wen, J., He, J., Tu, H., Kwak, Y., Eschstruth, A., Cayrel, A., Endre, G., Zhao, P.X., and Chabaud, M. (2008). Large_-_scale insertional mutagenesis using the *Tnt1* retrotransposon in the model legume *Medicago truncatula*. The Plant Journal 54, 335–347.

63. Thoquet, P., Ghérardi, M., Journet, E.-P., Kereszt, A., Ané, J.-M., Prosperi, J.-M., and Huguet, T. (2002). The molecular genetic linkage map of the model legume Medicago truncatula: an essential tool for comparative legume genomics and the isolation of agronomically important genes. BMC Plant Biology 2, 1–13.

64. Udvardi, M., Below, F.E., Castellano, M.J., Eagle, A.J., Giller, K.E., Ladha, J.K., Liu, X., Maaz, T.M., Nova-Franco, B., and Raghuram, N. (2021). A research road map for responsible use of agricultural nitrogen. Frontiers in Sustainable Food Systems 5.

65. Uffelmann, E., Huang, Q.Q., Munung, N.S., De Vries, J., Okada, Y., Martin, A.R., Martin, H.C., Lappalainen, T., and Posthuma, D. (2021). Genome-wide association studies. Nature Reviews Methods Primers 1, 59.

66. Villaécija-Aguilar, J.A., Hamon-Josse, M., Carbonnel, S., Kretschmar, A., Schmid, C., Dawid, C., Bennett, T., and Gutjahr, C. (2019). SMAX1/SMXL2 regulate root and root hair development downstream of KAI2-mediated signalling in Arabidopsis. PLoS genetics 15, e1008327.

67. Waidmann, S., Beziat, C., Ferreira Da Silva Santos, J., Feraru, E., Feraru, M.I., Sun, L., Noura, S., Boutte, Y., and Kleine-Vehn, J. (2023). Endoplasmic reticulum stress controls PIN-LIKES abundance and thereby growth adaptation. Proc Natl Acad Sci U S A 120, e2218865120.

68. Whitford, R., Fernandez, A., Tejos, R., Pérez, Amparo C., Kleine-Vehn, J., Vanneste, S., Drozdzecki, A., Leitner, J., Abas, L., Aerts, M., Hoogewijs, K., Baster, P., De Groodt, R., Lin, Y.-C., Storme, V., Van de Peer, Y., Beeckman, T., Madder, A., Devreese, B., Luschnig, C., Friml, J., and Hilson, P. (2012). GOLVEN secretory peptides regulate auxin carrier turnover during plant gravitropic responses. Developmental Cell 22, 678–685.

69. Wolner, B.S., and Gralla, J.D. (2001). TATA-flanking sequences influence the rate and stability of TATA-binding protein and TFIIB binding. Journal of Biological Chemistry 276, 6260–6266.

70. Xiao, T.T., Schilderink, S., Moling, S., Deinum, E.E., Kondorosi, E., Franssen, H., Kulikova, O., Niebel, A., and Bisseling, T. (2014). Fate map of Medicago truncatula root nodules. Development 141, 3517–3528.

71. Yoder, J.B., Stanton-Geddes, J., Zhou, P., Briskine, R., Young, N.D., and Tiffin, P. (2014). Genomic signature of adaptation to climate in Medicago truncatula. Genetics 196, 1263–1275.

72. Yoder, J.B., Briskine, R., Mudge, J., Farmer, A., Paape, T., Steele, K., Weiblen, G.D., Bharti, A.K., Zhou, P., and May, G.D. (2013). Phylogenetic signal variation in the genomes of Medicago (Fabaceae). Systematic biology 62, 424–438.

73. Yu, C., Miao, R., and Khanna, M. (2021). Maladaptation of US corn and soybeans to a changing climate. Scientific reports 11, 12351.

74. Zhang, J., Mazur, E., Balla, J., Gallei, M., Kalousek, P., Medveďová, Z., Li, Y., Wang, Y., Prát, T., Vasileva, M., Reinöhl, V., Procházka, S., Halouzka, R., Tarkowski, P., Luschnig, C., Brewer, P.B., and Friml, J. (2020). Strigolactones inhibit auxin feedback on PIN-dependent auxin transport canalization. Nature Communications 11, 3508.

